# In situ cryo-ET defines the ultrastructure of ER exit sites in human cells

**DOI:** 10.1101/2025.07.29.667472

**Authors:** Katie W Downes, Julia R Flood, Andrea Nans, Sander E. Van der Verren, Anjon Audhya, Giulia Zanetti

## Abstract

Trafficking of secretory proteins from the endoplasmic reticulum (ER) to the Golgi apparatus comprises the first, essential steps toward the appropriate localisation of 30% of eukaryotic proteins. Coat protein complexes COPII and COPI are involved in forward and retrograde transport of cargo and cargo receptors between the ER and the Golgi, respectively. Although COPII forms coated vesicles in vitro, the biogenesis, morphology and organisation of transport carriers in mammalian cells is debated. Here we use in situ cryo-electron tomography and super-resolution fluorescence microscopy to reveal the molecular architecture of ER exit sites in human cells that were not perturbed with drugs, temperature blocks, or overexpression systems. We visualise ribosome-exclusion zones enriched with COPII and COPI-coated vesicles and thus resolve the debate regarding the existence of COPII coated vesicles. COPII vesicles derive from ER membranes, whereas COPI vesicles originate from vesicular-tubular clusters that constitute the ER-Golgi intermediate compartment (ERGIC). We quantify coated vesicle morphology and positioning with respect to other ER exit site components, providing a molecular description of the organisation of the mammalian early secretory pathway.

## Introduction

The secretory pathway is responsible for the delivery of approximately 30% of eukaryotic proteins to their functional locations, including the majority of membrane and secreted proteins, and its components are essential and highly conserved^1–3^. Despite their importance, many of the molecular mechanisms that regulate protein secretion are poorly understood^4,5^.

The secretory pathway starts with the transport of newly synthesised proteins from the endoplasmic reticulum (ER) to the Golgi apparatus, where they acquire posttranslational modifications and are sorted. ER exit occurs at specific subdomains of the ER, called transitional ER, which in most organisms appear as punctate structures when fluorescently labelled via their stable components^6^.

In mammalian cells, where the Golgi is often concentrated in the perinuclear region, the ER-Golgi intermediate compartment (ERGIC) sits between sites of transitional ER and the cis-Golgi^6^. The transitional ER-ERGIC interfaces, commonly referred to as ER exit sites (ERES), are found across the whole cell volume and are characterised by ribosome-free areas enriched with vesicular tubular membranes^7^. ERES proximally positioned to cis-Golgi cisternae enable rapid cargo transit through the early secretory pathway, whereas cargo from peripheral ERES traverse the cytoplasm towards the Golgi via mobile ERGIC compartments^8,9^.

ER exit is dependent on a set of cytosolic proteins that together form the coat protein complex II (COPII)^5^. According to the prevalent model, born from early observations of membrane vesicles near the Golgi of secretory cells^10,11^ and supported by decades of biochemical and structural studies^12–20^, COPII assembles at transitional ER to form a coat that remodels the membrane into 60-100 nm vesicles, while simultaneously recruiting cargo proteins into the budding carriers. Upon uncoating, carriers fuse with target ERGIC membranes to release their cargo. Cargo receptors and adaptors, together with escaped ER resident proteins, are retrieved from either the ERGIC or cis-Golgi and transported back to the ER via retrograde vesicular transport mediated by the coat protein complex I (COPI)^21,22^, which is recruited to ERES and has been shown to act sequentially to COPII^23–25^.

COPII and COPI membrane budding has been reconstituted in vitro using purified coat components to reveal the molecular mechanisms of coat assembly and membrane remodelling ^13,16,26–28^. During COPII-mediated ER exit, the small GTPase Sar1 is activated by the ER-resident guanine nucleotide exchange factor (GEF), Sec12^29^. Upon activation, Sar1 stably inserts its amino-terminal amphipathic helix into the ER membrane and recruits the inner layer of the coat, comprised of Sec23-Sec24 heterodimers^30^. Sec24 is mainly responsible for recognition of cargo and cargo receptors^18^, while Sec23 recruits the outer layer of the COPII coat, made of rod-shaped Sec13-Sec31 heterotetramers that assemble to form cage-like structures^16,17,30^. Polymerisation of both the inner and outer coat contribute to generating and regulating membrane curvature^16^. COPII coated vesicles are transported to the ERGIC, in a manner regulated by condensates of TRK-fused gene protein (TFG), where they fuse to deliver their cargo^31^. Coat disassembly is facilitated by Sec23, which functions as a guanine nucleotide activating protein (GAP) for Sar1^30,32^, and is further stimulated by Sec31. The timing and regulation of the uncoating process remain unknown.

In the case of COPI, the small ribosylation factor Arf1 is activated by dedicated GEFs. Arf1 then inserts its myristoylated amino-terminus into Golgi or ERGIC membranes and recruits the 7-membered coatomer complex (formed by subunits 𝛼, 𝛽, 𝛽’, 𝛾, 𝛿, 𝜀, and 𝜁), en-bloc^21,22,33,34^. Coatomer polymerisation is coupled with the induction of membrane curvature, while cargo binding is orchestrated by the 𝛽 and 𝛿-COP subunits^21^. Two copies of ArfGAP are recruited by each coatomer, inducing stepwise GTP hydrolysis that initially enhances cargo recruitment and eventually leads to coat disassembly. Residual COPI coat subunits on the vesicle membrane might promote tethering to target membranes^21^.

Although COPII and COPI-coated vesicles have been visualised in situ by cryo-EM analysis of green algae^35^, direct evidence for this two-way vesicular transport system in vertebrates is lacking. Recent technological advances in high-resolution and live cell fluorescence microscopy have introduced doubts that the ‘classical’ vesicular model represents the main mechanism of COPII-mediated ER exit in higher eukaryotes. Live cell imaging in human cells showed that COPII puncta do not travel from their steady state location at ERES, whilst COPI is seen moving with ERGIC membranes to the Golgi^9,36^. Moreover, volume EM of ERES has shown what appear to be clusters of tubular vesicular membranes stably connected to the ER in COPII-positive regions, with COPI-positive tubules emanating from them, giving rise to the idea that COPII might form a collar to act as a ‘gate-keeper’ at tunnels directly connecting ER and ERGIC, rather than a coat at human ERES^8^.

The quest to discover ER exit mechanisms beyond the classic small vesicles model stems from the difficulty of explaining transport of the diverse range of cargo that are secreted in vertebrates, which include abundant and large proteins such as lipoproteins, collagens, and other extracellular matrix components^5,37,38^.

Here, we sought to map the molecular organisation of the early secretory pathway in human epithelial cells, with the aim to resolve controversies regarding the mechanisms of protein transport, and to provide a molecular-level understanding of the interplay between COPII, COPI, and other components of ERES.

We used cryo-correlative light and electron microscopy (cryo-CLEM) on unperturbed human retinal pigment epithelial (RPE-1) cells to target ERES for cryo-focused ion beam milling scanning electron microscopy (cryo-FIB/SEM) and acquire cryo-electron tomography (cryo-ET) datasets. We were able to resolve the COPII and COPI coats on membrane vesicles and buds. We also used stimulated emission depletion (STED) and confocal fluorescence imaging to quantify the relative positioning of ERES components. All together we provide a molecular description of ERES in human cells and our results support the vesicular transport model.

## Results

### Identification of ERES in situ

Typical mammalian cells contain a few hundred ERES, as seen by fluorescence microscopy (Extended Data Fig. 1a,b). Given the limited volume that can be imaged via cryo-ET of FIB-milled lamellae (typically below 0.5 µm^3^), the probability of encountering an ERES by random milling and acquisition is on the order of 1 in 100, effectively impairing our ability to unequivocally localise ERES and obtain non-ambiguous datasets containing our target of interest. To improve targeting of ERES in a live-cell compatible manner, we adopted a cryo-CLEM approach.

Making use of previously established human epithelial cells that express a functional Halo-tagged version of Sec23A at its endogenous locus^39^, we visualized HaloTag_Sec23A, via the Oregon Green Halo ligand, to direct the production of FIB-milled lamella and guide cryo-ET acquisition (Extended Data Fig. 1d). However, the presence of false positive autofluorescence puncta in cryo-conditions, together with difficulties in aligning the fluorescence data between the iFLM and TEM search maps and the lack of sign-posting features in the TEM, (Extended Data Fig. 2a,b,e,f) led to sites of interest being easily missed within the small field of view available with high magnification tomography (∼900 nm X 900 nm for a 2.24 Å pixel size).

To further improve targeting success, we developed a correlative approach where low-magnification, low-dose and high-defocus tomograms were used to select potential regions of interest (Extended Data Fig. 2c,d,g,h). Using this pipeline, we identified 134 regions of interest for high-magnification data collection, of which 28% clearly contained clusters of ER-derived coated vesicles and buds (Fig. 1). The remaining 72% contained membrane structures that did not appear coated or were not ER-associated and therefore were not analysed further.

**Figure 1.**
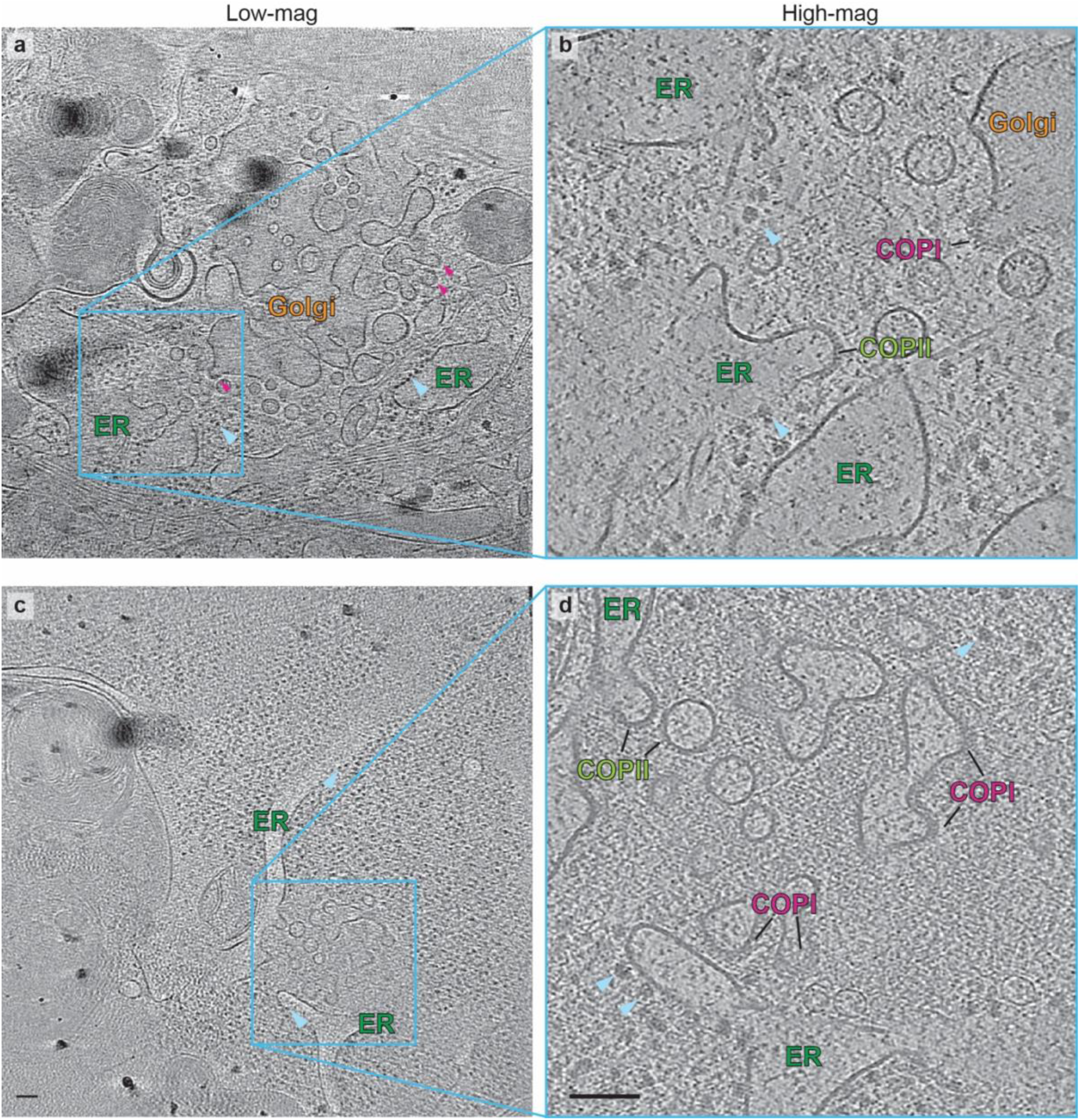
Cryo-ET of ERES in HaloTag-Sec23A RPE-1 cells. a. Slice through the xy plane of a reconstructed tomogram from a high-defocus, low-magnification tilt series from a FIB/SEM lamella of HaloTag-Sec23A RPE-1 cells, providing a panoramic view of cellular features. The ER is studded with ribosomes (pale blue arrowheads), while Golgi cisternae appear fenestrated and enriched with coated vesicles (purple arrowheads). b. Slice through the xy plane of a reconstructed tomogram from a high-magnification tilt series collected in the area highlighted in the cyan box in A, acquired at a resolution where molecular features such as COPII and COPI-coated membranes become apparent. c. As in A, but from a different lamella where no Golgi apparatus is visible. d. As in B. Scale bars 100 nm.

### Identification of COPII and COPI coats within ERES

ERES are known to be ribosome exclusion zones^7^. We defined ribosome exclusions zones based on 3D distribution maps of ribosomes, obtained by performing template-matching and subtomogram averaging of ribosome particles (Extended Data Fig. 3a,b). We picked over 16,000 ribosomes, and subjected them to subtomogram averaging following the WARP, relion, and M pipeline^40^ . This gave rise to a map at 15 Å resolution (Extended Data Fig. 3c-i), and to ribosome distribution coordinates that we used to quantify ribosome exclusion zones (Extended Data Fig. 4a-c). Further, we used ribosome subtomogram averaging to improve overall tilt series alignment and to determine the absolute hand of our reconstructions, validating an overall information content that extends to at least 15 Å.

Inspection of the cryo-tomograms allowed detection of vesicle coats within ribosome exclusion zones (Extended Data Fig. 4a). Comparison to coat density from previous in vitro reconstitution studies^16,28^ allowed COPII and COPI to be visually identified (Fig. 2a,b and Extended Data Fig. 5). Whereas COPI consists of a thick, fuzzy layer around the membrane (Fig. 2b, magenta boxes, Extended Data Fig. 5b), COPII has a well-defined, thinner inner coat layer, and a sparser outer layer (Fig. 2a, green boxes, Extended Data Fig. 5a). The outer layer was only visible in some slices in our in situ data, due to its long and thin structure which tends to be buried in the background of cytosolic components (Fig. 2a, red arrowheads).

**Figure 2.**
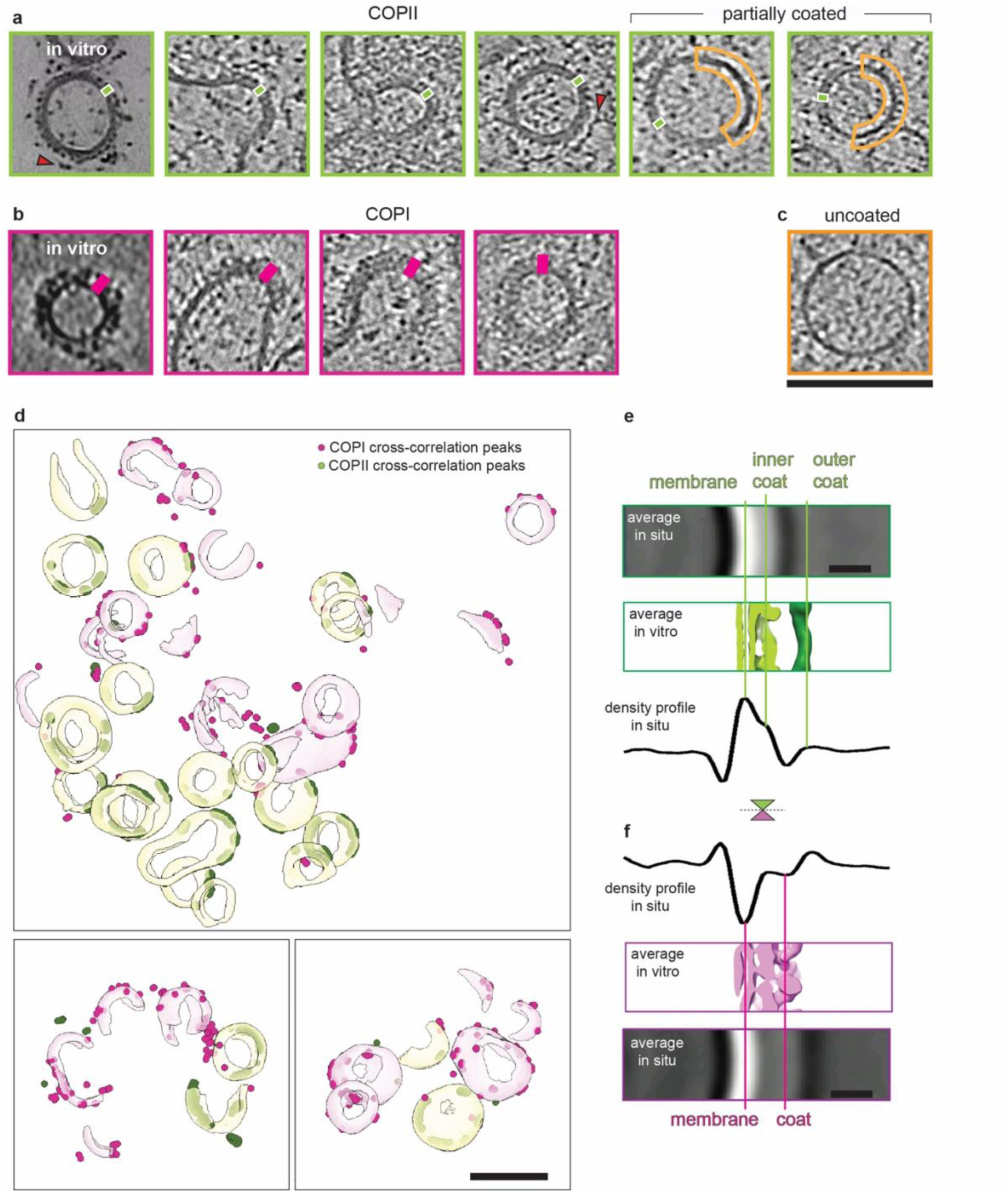
Identification of COPII and COPI-coated vesicles. a. Gallery of xy slices through COPII-coated vesicles from cryo-lamellae. Top-left panel shows an in vitro reconstituted vesicle for comparison^16^. Green rectangles are shown as rulers for coat thickness. Red arrowheads point to outer coat. The right panels show partially coated vesicles, with the uncoated region highlighted in yellow. b. Gallery of xy slices through COPI-coated vesicles from cryo-lamellae. Top-left panel shows an in vitro reconstituted vesicle for comparison^28^. Magenta rectangles are shown as rulers for coat thickness. c. An xy slice through an example of uncoated vesicle. d. 3D rendering of segmented coated membranes are shown in pale green and purple depending on whether they were manually assigned to COPII or COPI respectively. The highest cross-correlation peaks upon template matching using EMD-3968 as template (COPI structure) are shown as pink spheres, and those obtained with EMD-19417 as template (COPII structure) are shown in green. Template matching discriminates between the two classes of vesicles and there is very good agreement with our previous manual assignment. Three exemplary tomograms are shown. e. Top: Slice through the yz plane of the subtomogram average of particles picked from random oversampling of COPII-coated membranes, oriented normal to the membrane, and aligned via shifts only. Middle: the profile of an in vitro reconstituted sample is shown for comparison. Bottom, density profile obtained from the top panel. f. As in e, but for COPI. Panels are mirrored for clarity of comparison. Scale bars: a-d 100 nm (including all panels), e-f 10 nm.

We sought unbiased confirmation of our manual coat assignment via template matching. Using a semi-automated membrane segmentation routine, we created loose masks around all coated membranes (Extended Data Fig. 5d-i) and searched masked tomograms using available COPI or COPII structures as templates (EMD-3968 and EMD-19417, respectively). Both templates were run against identical masked tomograms. The two classes of coat were clearly discriminated, with the COPII template preferably correlating with coated membranes that we previously identified as COPII, and vice versa (Fig. 2d, Supplementary Video 1). The templates for both COPI and COPII were matched in positions and orientations that are consistent with the membrane curvature, further validating the peaks (Extended Data Fig. 5g,h). As a control, we performed template matching against a randomly masked tomogram which yielded disordered matched templates (Extended Data Fig. 5f,i), giving us confidence in our assignments.

We extracted COPI and COPII coat segments by randomly and evenly sampling corresponding coated membranes, assigned yaw and pitch angles normal to the membrane, and randomised their in-plane rotation. Upon 5 iterations of shift-only alignments and averaging we traced the density profile for each coat (Fig. 2e,f). For COPII, the profile was characterised by an intense double peak, and a weaker distal peak (Fig. 2e). This arrangement is consistent with the primary peak being formed by the membrane bilayer and the tightly juxtaposed inner coat, and the secondary peak resulting from the sparser outer coat, as reported previously in structures of in vitro reconstituted yeast proteins^15,16,41^ (Fig. 2e). A similar exercise repeated for COPI yielded an average profile consistent with the presence of a single ∼50 Å thick coat layer immediately juxtaposed to the membrane bilayer, as described in previous reconstitution experiments performed with mouse proteins and in situ studies of the green alga Chlamydomonas reinhardtii ^27,35^ (Fig. 2f).

### The Ultrastructure of ERES – COPII and COPI vesicles

With confidence in our assignment of coat identity, we were able to quantitatively characterize the molecular ultrastructure of ERES in 63 cryo-tomograms (Fig. 3 and Extended Data Fig. 6).

**Figure 3.**
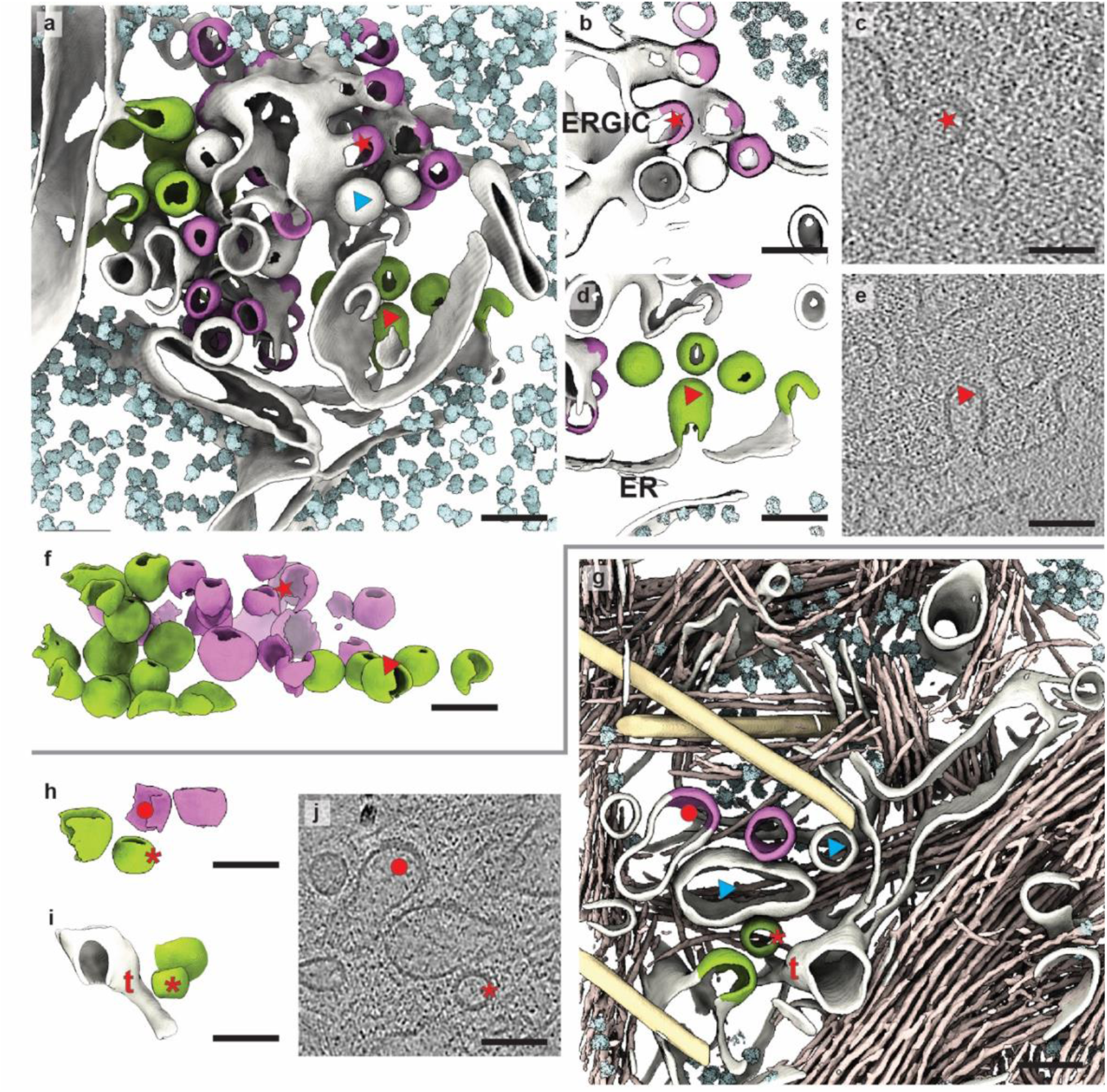
Molecular organisation of ERES. a. 3D rendered segmented tomogram of ERES, where membranes are white, COPII-coated membranes are green, COPI-coated membranes are purple, and ribosomes are pale cyan. b. A detail from the tomogram in a showing a vesicular-tubular cluster (ERGIC) with COPI buds and vesicles. c. An xy slice through the detail shown in b, cutting through a COPI coated bud (red star). d. A detail from the tomogram in a showing a vesicular-tubular cluster with COPII buds and vesicles. An ER cisterna, as identified by the presence of ribosomes on the membrane, is highlighted. e. An xy slice through the detail shown in d, cutting through a COPII coated bud (red triangle). f. COPII (green) and COPI (purple) segmented membranes are shown in isolation and from a side view, to display their vesicular nature and their size compare to the tomogram thickness. g. 3D rendered segmented tomogram of another ERES, where membranes are white, COPII-coated membranes are green, COPI-coated membranes are purple, ribosomes are pale cyan, intermediate filaments pale orange and microtubules pale yellow. h. as in f but for the tomograms shown in panel g i. a side view of a detail of the tomogram in g, directly comparing the appearance of COPII vesicle (red asterisk) with an ER tubule (red ‘t’). j. An xy slice through a detail of the tomogram in g, cutting through a COPII coated vesicle (red asterisk), a COPI coated bud (red circle). The red symbols mark the same structures across panels a-f. Cyan triangles in panels a and g mark uncoated vesicles and pleiomorphic structures. Scale bar 100 nm for all panels.

We found a total of 157 COPII coated membranes (95 vesicles and 62 buds), and 187 COPI coated membranes (39 vesicles and 148 buds). Coated membranes appeared clustered, occupying volumes with median diameters of 243nm (Inter Quartile Range (IQR): 168 to 371 nm), as measured by fitting an ellipsoid around all coated vesicles in each tomogram (Extended Data Fig. 4a,d).

Ribosome exclusion zones tended to loosely envelop coated vesicle areas, with diameters of 483 nm (IQR: 453 to 522 nm) (Extended Data Fig. 4d), and showed no obvious difference in electron density compared to the rest of the cytoplasm (Extended Data Fig. 3a). Ribosome density within ERES regions was on average 11.5 fold lower than in ribosome-occupied regions (Wilcoxon signed-rank test, n = 49, p = 3.6 × 10⁻¹⁵) (Extended Data Fig. 4e).

COPII coated membranes were observed either as membrane buds attached to the ER, or as free vesicles in close proximity to, but clearly separated from, the ER membrane (Fig. 3d,e and Extended Data Fig 6). We note that the thickness of our tomograms (150-350 nm) allowed for full vesicles to be resolved, leaving no ambiguity as to whether these are truly vesicles or sections through a tube (Fig. 3f,h,i). To confirm the displacement of COPII coated vesicles from the ER membrane we used confocal fluorescence microscopy, taking advantage of the scaffolding protein Sec16A as the marker for transitional ER. We imaged HaloTag-Sec23A RPE-1 cells stained with JFX650-HaloTag ligand and anti-Sec16 antibodies, which appeared only partially overlapping (Fig. 4a). Quantification of the displacement between transitional ER and COPII revealed a median distance of 106 nm within a 0 to 500 nm range (Fig. 4b, yellow).

**Figure 4.**
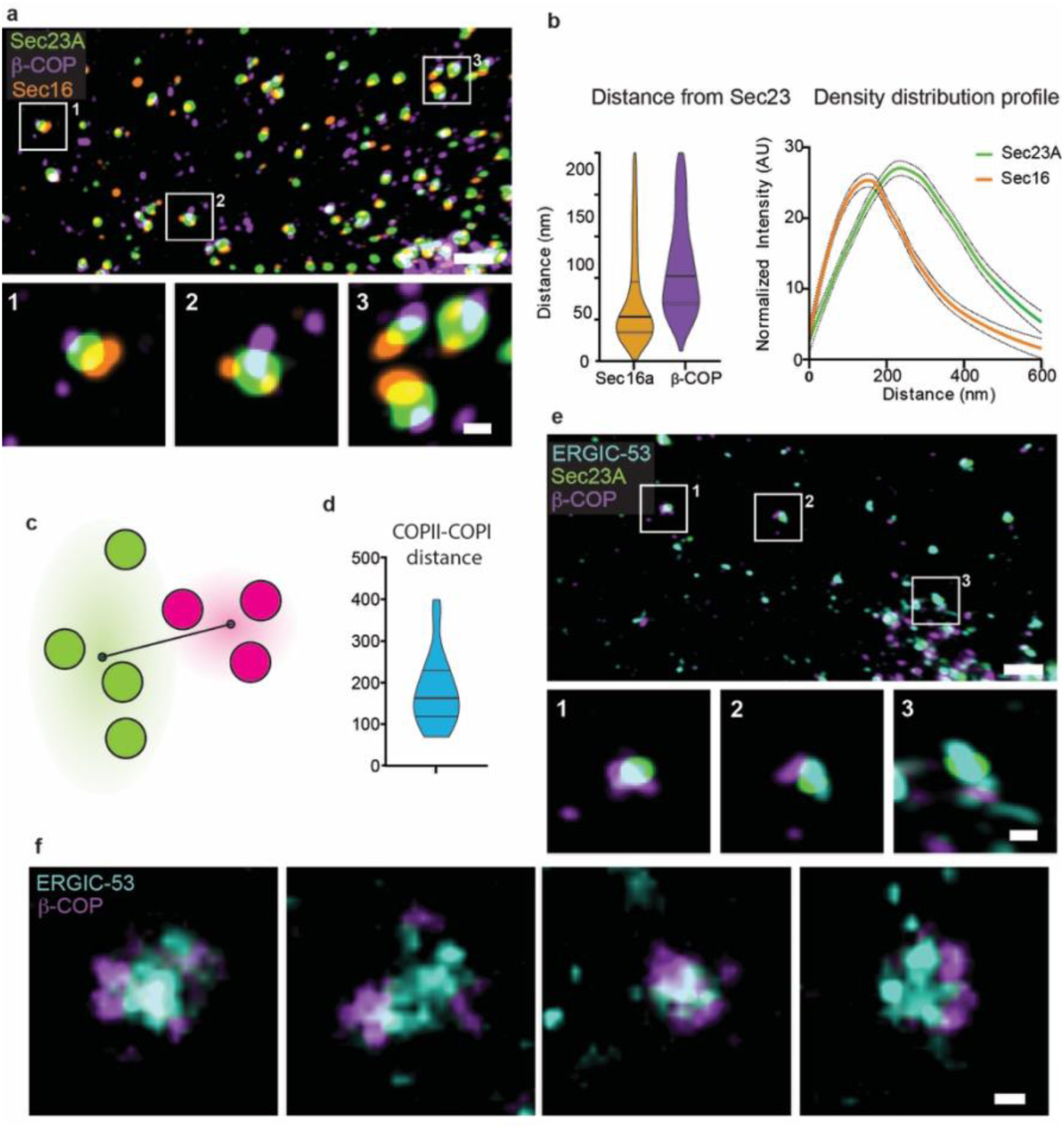
Fluorescence imaging of ERES components. a. Three-colour high resolution confocal fluorescence imaging of endogenous HaloTag-Sec23A RPE1 cells dyed with HaloTag JFX650 ligand (green) and stained with antibodies directed against Sec16 (orange) and 𝛽-COP (purple). b. Quantitative data highlighting the relative distribution of COPII (HaloTag-Sec23A) and transitional ER (Sec16A) and COPI (𝛽-COP). Left panel: Violin plots of the distance between centroids of endogenous HaloTag-Sec23A puncta from immunostained Sec16 and 𝛽-COP. Extracted from the three-color high resolution confocal fluorescence imaging as exampled in A. For Sec16, n = 7666, from ≥10 cells, across 3 experimental repeats. For 𝛽-COP, n = 1301, from ≥10 cells, across 4 experimental repeats. The lines represent the median and IQR: Sec16 median and IQR: 106, 68.7-190 nm. COPI median and IQR: 204, 138-298 nm. Right panel: Averaged linescans of Sec16 (orange) and HaloTag-Sec23A (green), extracted from two-colour super resolution STED imaging and fitted from 50 structures across 2 cells over 1 experimental repeat. Standard error is shown by the dotted lines. The intensity values were normalised for display purposes. c. A schematic summarising how distances between the centre of mass of COPII and COPI vesicle clusters were measured on tomograms, as plotted in panel e. d. Quantitation of the distances between COP2 and COP1 clusters, measured as described in Figure 4d from all ERES containing tomograms. Median = 163 nm. IQR = 118 – 229 nm. e. Three-color high resolution confocal fluorescence imaging of endogenous HaloTag-Sec23A RPE1 cells dyed with HaloTag JFX650 ligand (green) and stained with antibodies directed against ERGIC-53 (cyan) and 𝛽-COP (purple). f. Two-colour super resolution STED imaging of endogenous SNAP-tag-ERGIC-53 RPE1 cells stained with antibodies directed against 𝛽-COP (purple) and SNAP-tag-ERGIC-53 dyed with JFX650-SNAP-tag ligand (cyan). n=1 Scale bar: A,C overviews: 1 µm, insets: 200 nm, D: 100 nm Experiments in panels a and e were repeated a minimum of two times. Experiment in panel f was repeated once. Source numerical data are available in source data

COPI coated structures were found in proximity to COPII coated membranes in 86% of the tomograms. In those tomograms where COPI is not visible, we could not establish whether COPI was present in the vicinity of COPII but was excluded from the field of view, or truly absent. Using 3-color confocal imaging, we found COPI near almost all Sec23A puncta (Extended Data Fig. 7a), at a median distance of 204 nm (IQR: 138 to 298 nm) (Fig. 4b, purple) and often on the distal side from Sec16A (Fig. 4a). The distance between clusters of COPII and COPI vesicles measured in the tomograms was 163 nm (IQR: 118 to 229 nm), which is in a range compatible with measurements from fluorescence (Fig. 4 c,d). 41% of COPI puncta were not associated with COPII puncta and most likely represent Golgi-derived COPI vesicle clusters (Extended Data Fig. 7a). We note that many tomograms in the full cryo-ET dataset contained COPI but not COPII coated vesicles, however, we did not include these in our analysis as there is no evidence that these vesicles are associated with ERES.

Unlike COPII, ERES-localised COPI coated buds extended from membranous vesicular-tubular clusters distinct from the ER (Fig. 3a-c,g,j). Previous studies have shown the co-localisation of COPI and the ERGIC marker ERGIC-53^42^. We used confocal fluorescence imaging to assess the relative position of COPI, COPII and ERGIC-53 in our cells, confirming the tight juxtaposition of both COPI and COPII with ERGIC-53 (Fig. 4e, Extended Data Fig. 7a). Super resolution 2-color STED of endogenously tagged ERGIC-53 RPE-1 cells further showed that COPI vesicles were tightly co-localised with ERGIC-53 puncta and often appeared to encircle them (Fig. 4f). These data strongly suggest that the compartments from which we observed COPI vesicles originate are ERGIC membranes.

Most ERES had between 1 and 3 COPII coated membranes, inclusive of buds and vesicles (Fig. 5), however we observed a few instances of ERES with up to 19 COPII events (Fig. 5a). The number of COPI events in the majority of ERES was also between 1 and 3, with exceptional cases of up to 21 (Fig. 5b). We note this is likely an underestimation as the overall size of an ERES in 3 dimensions will often exceed the thickness of the tomogram by 2-3 fold. On average, from our ERES-containing tomograms, there were more COPI than COPII events per ERES. However, due to the proximity of the Golgi in several tomograms, it was often impossible to know whether COPI vesicles were ERGIC or Golgi-derived, leading to a potential overestimation of their number at ERES. Nevertheless, we detected a significant positive correlation between the number of COPII and COPI events at each ERES (R=0.53, P=1.5E-9), indicating their functional and/or morphogenetical interdependence (Fig. 5c).

**Figure 5.**
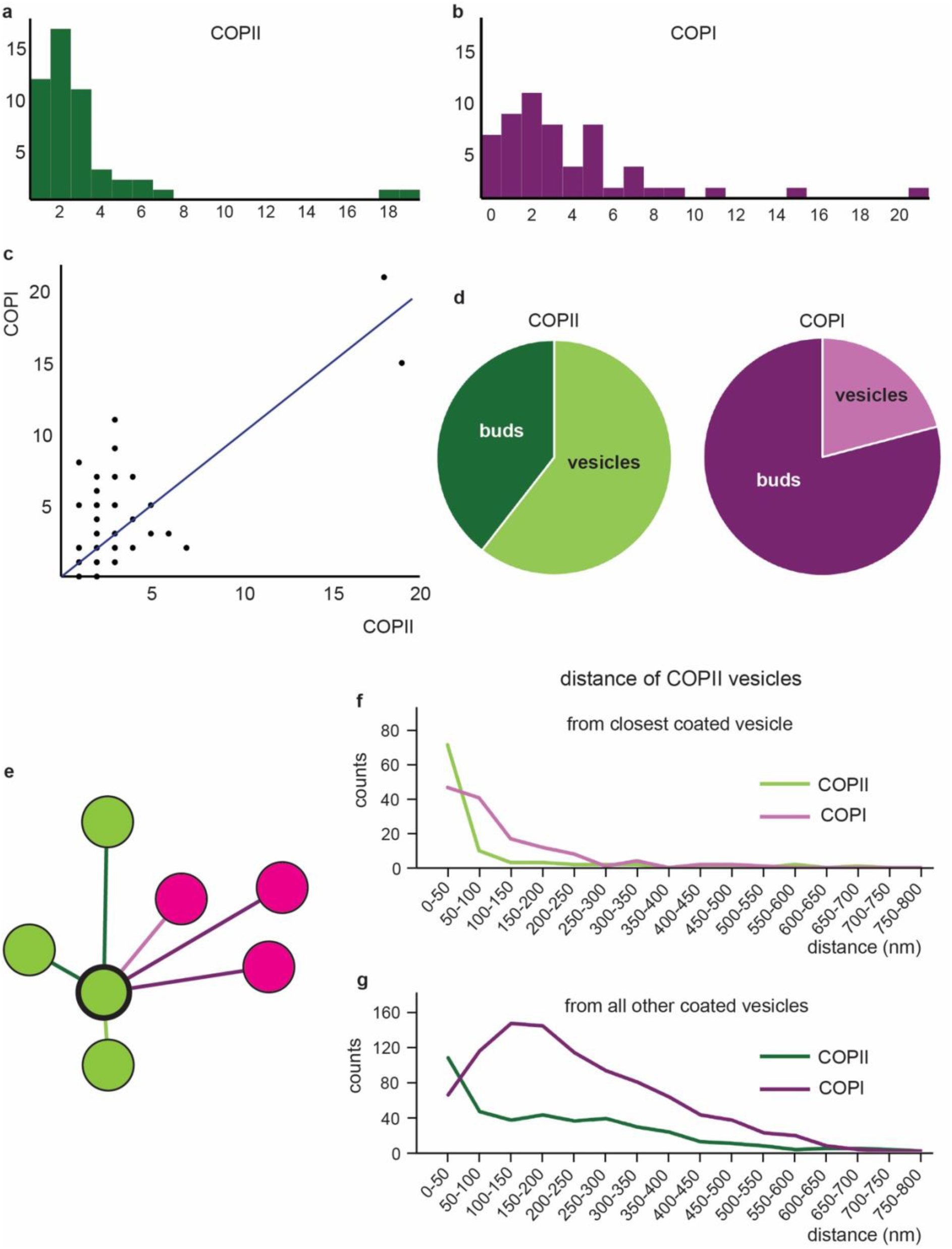
Distribution of coated vesicles at ERES. a. Distribution of the number of COPII-coated events per ERES (n=157). b. Distribution of the number of COPI-coated events per ERES (n=187). c. Scatter plot of the number of COPI vs COPII coated events. The blue solid line represents x=y. d. Pie chart showing the relative abundance of buds vs vesicles for COPII (green) and COPI (purple). e. schematic to illustrate the distance analysis performed: we measured distances between the closest membrane points of each pair of COPII vesicles (green lines) and of COPII-COPI (purple lines). The closest pair is depicted in lighter coloured lines. f. Distribution of distances of all COPII vesicles from their nearest COPII neighbour (green line) and COPI neighbour (pink line). g. Distribution of distances of all COPII vesicles from all COPII neighbours in the same tomogram (dark green line) and all COPI neighbours in the same tomogram (purple line). Source numerical data are available in source data

To further probe the relative positions of COPII and COPI, we measured membrane-membrane distances of all pairs of coated membranes (Fig. 5e). When we assessed the distribution of distances of all COPII vesicles or buds from their coated neighbours, we found that nearly all COPII coated membranes had their nearest COPII neighbours within a 100 nm range (Fig. 5f, green line), while their nearest COPI neighbours tended to be further away (Fig. 5f, pink line). The distribution of distances between all pairs of COPII membranes showed a steadily declining profile (Fig. 5g, dark green line), indicative of COPII buds and vesicles tendency to cluster. The distribution of distances between all pairs of COPII and COPI membranes showed a marked peak between 100 and 200 nm, with fewer COPI membranes being closer to COPII than 100 nm (Fig. 5g, purple line), suggesting some level of spatial separation between the two types of vesicles.

Non-coated vesicles and non-coated pleomorphic structures were also observed within the ribosome exclusion zone and near both COPII and COPI vesicles (Fig. 3a,g, cyan triangles, Fig. 2c). Occasionally, partially coated COPII vesicles were seen, corroborating the hypothesis that uncoating happens before fusion (Fig. 2a, right panels). However, such events were hard to quantify due to the high noise levels associated with in situ data, the fact that some vesicles were only partially contained in the tomogram volume, and the anisotropy of the reconstruction that weakens features along the view axis. Whether uncoated membranes are mechanistically involved in ERES function is unknown, but it is plausible they represent previously coated structures, which act as fusion competent intermediates.

### Transport Carrier Morphology at ERES

As described above, we detected fully detached coated vesicles as well as coated membrane buds (Fig. 3 and Fig. 5). To confirm coated vesicles are not sections through tubes, we fitted ellipses to axial 2D projections of segmented vesicles and calculated their circularity as the ratio between the shortest and longest elliptical axis. Closed vesicles will have values close to 1 while projections of randomly oriented tubes are expected to range from spherical to strongly elliptical. Both coated vesicles ranged from ∼0.8 – 1 with a median of ∼0.9 (Extended Data Fig. 7b) showing that both types of coated vesicles tend to be close-to-spherical in shape.

COPII buds were always associated with ER membranes, whereas COPI buds were always associated with ERGIC or Golgi membranes (Fig. 3 and Extended Data Fig. 6). This suggests that, for both COPII and COPI, we detected budding, rather than fusion events. We never saw uncoated buds associated with either ER or ERGIC membranes, indicating that for both coats, uncoating occurs after scission, but before fusion with the target compartment, and that fusion of transport vesicles is rapid and unlikely to be detected in tomograms. COPII appeared to be more often associated with detached free vesicles than COPI (Fig. 5d), suggesting scission and uncoating occur over distinct timescales for the two coats.

We performed morphometric analysis^43^ to fit the segmented and extracted coated membranes to ellipsoids or spheres and quantify their volume (Extended Data Fig. 7c). COPII coated vesicles had a median volume of 180,760 nm^3^ (Extended Data Fig. 7d), which when approximated to a sphere corresponds to a diameter of 70 nm, consistent with previous reports^12,13,16^. The COPII coated vesicle volumes ranged from 78,007 nm^3^ to ∼400,000 nm^3^ (diameters of 53.3 nm and 91 nm, Extended Data Fig. 7d).

COPI coated vesicles had a significantly smaller median volume of 134,153 nm^3^ (diameter of 64 nm, P=0.0014, Extended Data Fig. 7d). This is slightly larger than previously reported for both in situ Chlamydomonas vesicles and vesicles reconstituted in vitro with mouse proteins, which have mean diameter of ∼55 nm^28,35^. However, this previously reported volume is included within the range observed in this dataset (Extended Data Fig. 7d), which corresponds to a spherical diameter of 47 nm to 131 nm. Our measurements nevertheless suggest there may be morphological differences in COPI coated vesicles across evolution.

The distributions of the volume of COPI and COPII vesicles were also significantly different (P=0.0014), with COPI displaying a greater overall range and a pronounced positive tail (Extended Data Fig. 7d). This suggests that the COPI machinery may tolerate a greater range in carrier size whereas COPII may allow less flexibility.

Morphometric analysis of coated membrane buds still attached to the donor compartment was more challenging than vesicles, due to the range of morphologies present in the data and the presence of an ‘open face’. Nevertheless, ellipsoids could still be used to approximate their encompassed volume (Extended Data Fig. 7c). To this end the 149 COPI and 56 COPII coated buds present in our data were extracted and analysed. COPII coated buds enclosed volumes ranging from 32,229 nm^3^ to 782,226 nm^3^ with a median of 158,886 nm^3^ (diameter of 39 nm, 144 nm and 67 nm, Extended Data Fig. 7e). COPI coated buds were not significantly different than COPII in terms of both scale and distribution and ranged from 5,560 nm^3^ to 3,215,442 nm^3^ with a median of 141,587 nm^3^ (diameter of 22 nm, 183 nm and 65 nm, Extended Data Fig. 7e).

### The Ultrastructure of ERES – cellular context

Analysis of pairs of high and low-magnification tomograms provided detailed intracellular context. Golgi membranes could be identified in both high and low-magnification tomograms due to their stacked and fenestrated appearance (Fig. 1a,b and Extended Data Fig. 8) and thus the spatial relationship between ERES, ERGIC and Golgi could be extracted. In some cases, ERES were observed in proximity to Golgi membranes allowing their classification as perinuclear ERES, whilst others were distant from any visible Golgi (Fig. 1a,c and Extended Data Fig. 8). The latter could represent perinuclear sites where Golgi was removed by FIB milling, however proximity to the plasma membrane allowed us to characterize at least some of these as peripheral ERES (Extended Data Fig. 8b). ERGIC membranes, characterised by their vesicular tubular morphology and COPI-coated buds, were observed indiscriminately at both peripheral and centrally located ERES (Extended Data Fig. 8). When perinuclear ERES were only a few hundred nanometers away from Golgi membranes (Extended Data Fig. 8a), it could be hard to distinguish ERGIC from cis-Golgi. However, such distinction might be a matter of definition in a context where the ERGIC matures to become cis-Golgi.

It is unclear whether coated vesicles and ERGIC membranes rely on cytoskeletal filaments for their directed movement. We analysed our tomograms for the presence of filaments and found a great degree of heterogeneity (Fig. 3a,g). 72% of ERES tomograms contained microtubules (numbering between 1 and 4), but these didn’t appear to be preferentially associated with any of the membranes involved. 84% contained intermediate filaments, often forming an extended meshwork encompassing the entire ERES area. Finally, actin was observed in 17% of tomograms. In all cases, no significant difference was found in the frequency of filaments in tomograms with or without ERES (P=0.825 for microtubules, P=0.317 for intermediate filaments and P=1 for actin filaments), and thus a role for filaments specific to ERES cannot be envisaged or established from our data.

### TFG tethers COPII vesicles at ERES

TFG has been shown to cluster COPII transport intermediates at ERES, contributing to their uncoating via competition with outer coat component Sec31^44–47^. Confocal fluorescence imaging confirmed that TFG co-localised with HaloTag-Sec23A, and closely juxtaposed to other ERES components, including Sec12 (Fig. 6a). Recent studies suggest that TFG condensates may act as a sieve, enabling entry of COPII coat proteins but not assembled COPI^48^. However, the partially interspersed distributions of COPII- and COPI-coated membranes seen by cryo-ET (Fig. 5) is inconsistent with the existence of a TFG barrier separating the two classes of vesicles. Consequently, we sought to understand the spatial relationship between COPII coated vesicles and TFG. Due to the lack of recognizable features in cryo-tomograms to identify TFG, we used super resolution STED imaging (Fig. 6b). TFG appeared to be surrounded by COPII-positive membranes (Fig. 6c), consistent with TFG clustering COPII carriers, as opposed to restricting the entry of COPI.

**Figure 6.**
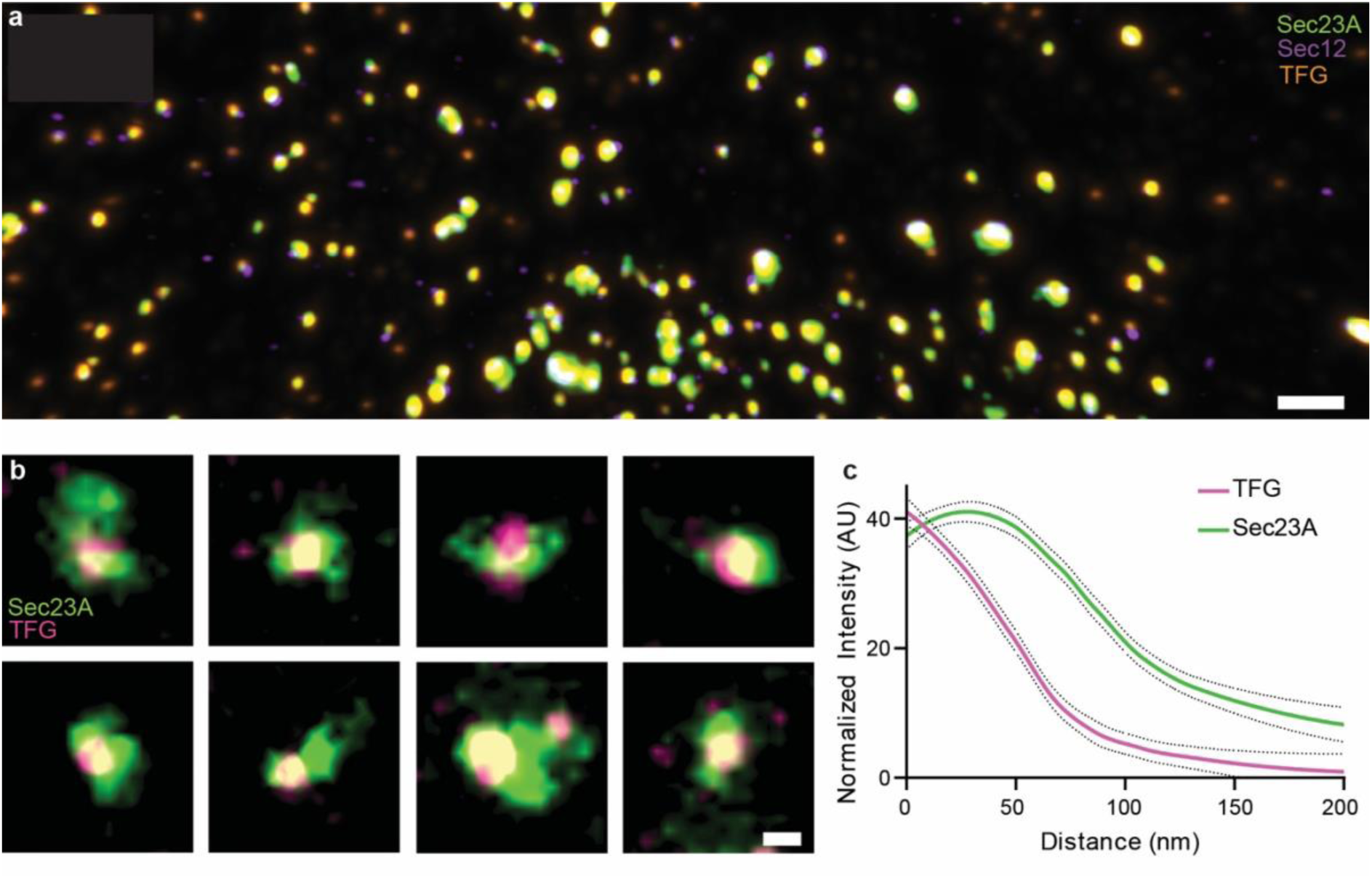
Co-localisation of TFG and Sec23A in HaloTag-Sec23A RPE1 cells. 1. Three-color high resolution confocal fluorescence imaging of endogenous HaloTag-Sec23A dyed with HaloTag JFX650 ligand (green) and immunostained Sec12 (purple) and TFG (orange). 2. Two-color super resolution STED imaging of endogenous HaloTag-Sec23A dyed with HaloTag JFX650 ligand (green) and immunostained TFG (pink). 3. Averaged linescans of TFG (pink) and HaloTag-Sec23A (green) fitted from 50 measurements across 9 cells over 3 experimental repeats, showing the distribution of HaloTag-Sec23A signal from the center of TFG condensates extracted from the two-color super resolution STED imaging. Standard error is shown with by the dotted lines. Scale bars overviews: 1 µm, insets: 100 nm

## Discussion

We used fluorescence and cryo-ET to visualize ERES in situ in a human cell line. We obtained cryo-tomograms with information extending to 1.5 nm resolution, where we could detect coated vesicles and define their molecular identity. We found that ERES consist of COPII and COPI coated vesicles, COPII coated buds attached to the ER, and vesicular-tubular membranes that are independent from the ER and that have COPI coated membrane buds. Using STED and confocal imaging, we defined these structures as ERGIC membranes.

In contrast to previous interpretations of lower-resolution fluorescence and volume EM data^8,36^, we did not observe tunnels connecting the ER with either ERGIC or Golgi membranes, nor did we find any evidence of “COPII-coated collars”. We were able to resolve several membrane tubules extending from the ER (Fig. 3i); however, they were not coated and always connected with another ER cisterna whenever we could trace their destinations, consistent with the well-established network-like nature of this organelle.

Our data do not exclude that COPII-derived tunnels exist in cells, as our workflow was biased towards clusters of vesicles. However, our results show that the main mode of ER exit – at least in unperturbed human epithelial cells – relies on COPII coated vesicles (Fig. 7), compatible with current understanding of coat-mediated membrane remodelling. The alternative scenario, where a COPII collar would serve as gatekeeper without forming a coat, currently lacks both experimental support and a physical mechanism for curvature generation.

**Figure 7.**
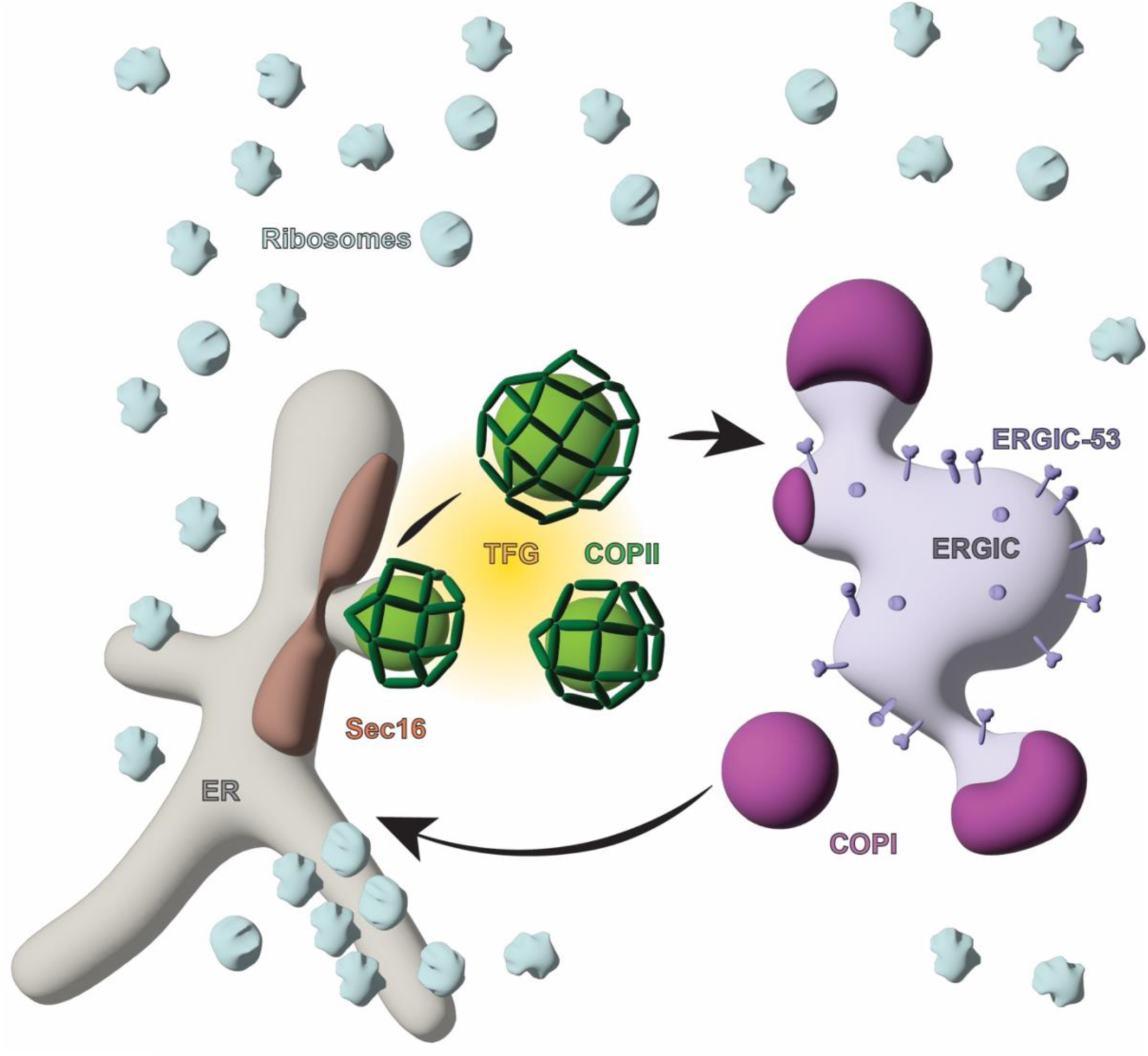
An updated model of the mammalian ERES. COPII coated vesicles bud off Sec16A-positive segments of the ER membrane in ribosome-free regions of the cytosol. COPII vesicles are clustered by TFG in the area between the transitional ER and the ERGIC. COPI vesicles bud off ERGIC vesicular-tubular membranes and are found in close proximity to COPII.

The molecular underpinning of how and when cargo is transported from ERGIC to Golgi remains to be resolved. Live cell fluorescence suggest microtubule motor-dependent movement, and it has been proposed that COPI might serve as an anterograde coat from the ERGIC to the Golgi^8^. It is unlikely that independent small COPI coated vesicles traverse the cytosol across tens of microns to be delivered to the Golgi. We believe our results are more compatible with a scenario where ERGIC clusters move en bloc towards the Golgi along microtubules, while COPI dynamically associates with the ERGIC to retrieve cargo receptors and escaped ER-resident.

While we did see microtubules in >70% of our tomograms, these did not appear to be associated with ERGIC membranes or COPI coated vesicles, and their numbers were similar to those seen in non-ERES containing tomograms. We hypothesize that ERGIC membranes might engage with microtubules at a temporally delayed point, after dissociation or dissolution of the source ERES. However, there is evidence that ERES are long lived^49^, so an alternative hypothesis is that anterograde ERGIC subdomains may detach from their source ERES and engage with microtubules at a spatially separated site. More work including live fluorescence and cryo-ET is needed to establish the relative lifetime and movement of ERES and ERGIC membranes and their cytoskeleton engagement.

Both COPI and COPII coated membranes in our tomograms had density features consistent with in vitro reconstituted vesicles, validating previous structural efforts to characterize their molecular architecture^15,16,28^. Coated vesicles and buds displayed a variety of shapes and sizes (Fig. 2, Fig. 3, Extended Data Fig. 5 and 7), consistent with flexible assembly mechanisms that provide a suitable platform for regulation. While our in situ data were not conducive to structure determination, the variability in size and shape of COPII coated carriers strongly suggests that the mode of assembly previously characterised in vitro, with locally ordered inner coat patches stabilised by outer cages of variable geometries^16^, is plausible.

Morphometric analysis revealed that COPI vesicles were on average smaller and more variable in size and shape than COPII (Extended Data Fig. 7d), potentially suggesting greater flexibility in the geometry of the COPI coat or less stringent regulation of vesicle scission.

Comparing the number of coated buds and free coated vesicles, we found that COPII and COPI displayed opposite behaviour (Fig. 5d). COPI was found more often around buds, indicative of either faster uncoating or slower curvature acquisition and delayed scission. GTP hydrolysis on Arf1 is triggered in stages by Arf-GAPs that are bound to different subunits of coatomer. It was proposed that the second wave of GTP hydrolysis, leading to uncoating, might happen once the COPI subunits are arranged around high membrane curvature (i.e. a complete vesicle)^21,27^. This would explain why we see few coated free vesicles, as uncoating would happen fast after vesicle completion.

In the case of COPII, the coat seemed more persistent on free vesicles (Fig. 5d). While the timing of COPII uncoating remains unclear, it has been shown that upon GTP hydrolysis of Sar1 the inner coat components persist for some time^50^, probably via interaction with cargo and maintenance of a stable outer cage. Full uncoating is likely to occur once the outer coat cage disassembles, weakening inner coat lattice arrangement. TFG has been implicated as an uncoating factor that acts by competing with Sec31 for Sec23 binding, and is a likely candidate for triggering full uncoating^44,51^. Our STED data suggests that TFG clusters neighbouring vesicles, consistent with a model where fully coated vesicles encounter TFG foci, where they are stripped down^44,51^.

Taking advantage of the individual vesicle resolution of cryo-ET, we analysed the distributions and organisation of COPII and COPI membranes. COPII vesicles are predominantly located within 100 nm of another COPII vesicle (Fig. 5f), and the probability of finding other COPII vesicles declines with increasing distances (Fig. 5g). This is a profile characteristic of clustering, likely reflecting COPII vesicle’s shared site of origin at the transitional ER. In contrast, the proximity of COPI vesicle to COPII is reduced (Fig. 5f), with the COPII-COPI distance distribution showing a distinct peak between 100 and 200 nm, and a notable scarcity of COPI vesicles within 100 nm of COPII (Fig. 5g). These data suggest that COPI and COPII vesicles maintain distinct albeit partially overlapping spatial domains.

Given that both vesicle populations traverse the same ER-to-ERGIC interface, it can be hypothesised that uncoating occurs before vesicles disperse to the point of completely mixing. In this context TFG could play a dual role of slowing down diffusion of COPII vesicles while aiding their uncoating^51^. Alternatively, COPI and COPII vesicles could travel along preferential routes, however the spatial distribution of the two types of vesicles, while distinct, has broad overlap (Fig. 5 f,g), suggesting the latter hypothesis is less likely. We saw no evidence supporting the proposed model for TFG to form sieve-like structures separating areas of COPII- and COPI-mediated transport^48^.

This work provides a framework to advance the ongoing discourse around the molecular nature of ER-derived transport carriers, but it is by no means exhaustive. COPII is required for ER exit of a variety of cargo in all cell types, and our results leave the open question of how ERES are uniquely organised in various specialised cell. Highly secretory cells tend to have significantly increased expression levels of secretory pathway components, including COPII. It is often the case that a particular paralogue of COPII is upregulated in such cells, for example Sar1B over Sar1A in plasma cells or early spermatids, or Sec24D in many secretory cells including plasmablasts and hepatocytes (https://www.proteinatlas.org/). It remains to be answered whether different paralogues influence the organisation of ERES and the morphology of carriers.

The organisation of the early secretory pathway in neurons will also need to be investigated. Although the majority of COPII-mediated transport has been suggested to take place in the cell body, ERES components are found in both axons and dendrites, potentially contributing to local expression of membrane proteins, including signalling receptors and cell adhesion factors^52^. Depletion or overexpression of a dominant negative isoform of Sar1 has been shown to inhibit axon outgrowth^53^. Additional cryo-ET studies are necessary to determine the structure and organisation of neuronal ERES.

It is also hard to reconcile what we observe in epithelial cells with ER exit of large cargo, which are physically incapable of being packaged into 60-90 nm transport intermediates and yet show dependence on COPII. Collagen secretion has long intrigued researchers as procollagens constitute one of the most abundant classes of secretory cargo, yet their size (up to 500 nm) is prohibitively large for incorporation into canonical vesicles. Several models for collagen ER exit have been proposed, from large-coated carriers to tubules and tunnels^37,38,54–59^. A number of factors that contribute to COPII-mediated transport are necessary for collagen secretion, and certain mutations in COPII only affect collagens, leading to disease^1,3,60–64^. It is therefore possible that COPII transport has adapted and evolved in collagen secreting cells. We speculate that regulation of ‘normal’ COPII functions could lead to the generation of procollagen competent carriers. For example, tuning of coat assembly and disassembly rates could be sufficient in producing larger or tubular carriers, and a delay in scission could lead to the generation of ‘tunnels’ connecting ER and ERGIC without re-inventing COPII molecular mechanisms. Factors from the Tango1 family are likely to be involved in this regulation^57,61^. The only way to answer this mystery is to directly visualize collagen carriers at a resolution sufficient to discern coated membranes.

## Acknowledgements

We thank the Structural Biology Science Technology Platform at the Crick for technical and computational support. We thank Elizabeth Miller from the University of Dundee and Patrick Phillips from the Francis Crick Institute for feedback on the manuscript. This work was supported by grants to G.Z. from the European Research Council (ERC-StG-2019 grant 852915) and UKRI (MR/Z504312/1). Work in the group of A.A. was supported in part by a grant from the National Institutes of Health (GM134865).

## Contributions

Conceptualisation, G.Z.; funding acquisition, G.Z., A.A; fluorescence microscopy, J.F and A.A. sample preparation for cryo-EM, K.D. and S.V.D.V.; cryo-FIB/SEM and cryo-ET data collection, K.D. and A.N.; cryo-ET data processing, K.D. and G.Z.; writing (original draft), K.D. and G.Z.; writing (revisions), all authors.

## Materials and Methods

This research complies with all relevant ethical regulations.

### Cell culture

Wild type human retinal pigmented epithelial cells (RPE-1) were obtained from ATCC and were cultured in DMEM/F12 (ThermoFisher Scientific) plus 10% FBS, L-glutamine and 1% Pen/Strep (Bio Basic). CRISPR modified Halo-Sec23A and SNAP-tag-ERGIC-53 RPE-1 cells were sourced as described in^39^ and maintained in analogous conditions to wild type. Briefly, CRISPR-Cas9 editing was used to create a translational fusion between HaloTag and the dominant Sec23 paralog, SEC23A, at its endogenous locus in RPE-1 cells, resulting in a homozygously edited line (Extended Data Fig.1c). Depletion of Sec23B in the edited cell line did not impair growth nor viability, and movement of secretory cargo was unaffected, indicating the tagged Sec23A is functional^39^.

### Fluorescence microscopy

For immunofluorescence studies, cells were labelled with JFX650-HaloTag ligand (generously provided by Luke Lavis) and subsequently fixed using 4% paraformaldehyde at 37°C for 8 minutes or 100% methanol at -20°C for 10 minutes, followed by permeabilisation and antibody labelling (1 mg/ml) at 4°C overnight. Immunofluorescence studies were conducted using validated antibodies against Sec16A (Bethyl Laboratories; A300-648A), Sec31A (BD Sciences; 612351), TFG (Novus Biologicals; NBP2-62212), ERGIC-53 (Santa Cruz Biotechnology; sc-66880), and COPB (Santa Cruz Biotechnology; sc-393615). After thorough washing, coverslips were incubated with secondary antibodies (AlexaFluor conjugates) for 1 hour at room temperature in the dark and mounted using Prolong Diamond Antifade. Slides were cured for 24 hours in the dark at room temperature before imaging.

Confocal imaging was conducted using a Nikon Ti2 spinning disk confocal microscope equipped with a 60x oil immersion objective (1.4 NA) and a Hamamatsu ORCA-Flash4.0 sCMOS camera. Imaging datasets were comprised of 25-35 sections separated by 0.3 µm (depending on sample thickness).

3-color confocal imaging was conducted on a Nikon AX R imaging system with NSPARC using a 100x SR HP Apo TIRF oil immersion objective (1.49 NA). The NSPARC detector allows for improved lateral resolution and signal to noise ratio. The TIRF objective allows more light to be collected and thus better resolution. Data were further processed by Nikon proprietary ‘denoise’ software settings to elucidate their geometry. Imaging datasets were comprised of 32-43 sections separated by 0.08 µm. The TetraSpeck Fluorescent Microspheres Size Kit (ThermoFischer) was imaged to establish the field of view without chromatic aberration, as determined by complete overlap of fluorescent beads in all channels. The center third of the field of view was identified as devoid of chromatic aberration and thus used for analysis.

For STED microscopy, cells were labeled with JFX650-HaloTag Ligand and fixed as described above. Alexa Fluor 594 secondary conjugates were used prior to mounting using Prolong Diamond Antifade. Coverslips were cured at room temperature in the dark for 24 hours prior to imaging on an Abberior STEDYCON super-resolution imaging system using the 775 nm depletion laser and a 100X Plan Apo Lambda objective (1.45 NA). This system does not require manual alignment and instead is equipped with an automatic alignment procedure.

### Confocal and STED Image Analysis

To identify structures, the Spots Creation Wizard was used in Imaris Microscopy Image Analysis Software (Oxford Instruments). The software subtracts background and identifies spots based on the intensity at the center of each spot. Quality thresholds were established per channel per experiment to eliminate background staining.

To quantify the number of ERES per cell, the total number of Sec31 positive punctate structures was counted in control RPE-1 cells (n = 32 cells, N = 2).

To quantify distance between markers in images acquired on the Nikon AXR system, distance thresholds were set to eliminate non-co-located structures. The threshold for HaloTag-Sec23A and Sec16A and HaloTag-Sec23A and β-COP was 0.5 µm. Various thresholds were tested to ensure data integrity. The distance between the centre intensities of co-located structures was quantified using the Shortest Distance tool in Imaris. For each condition at least 1900 distances were measured from at least 10 cells across 2 experimental repeats.

To quantify the average percentage colocalisation between COPI and COPII, a distance threshold of 0.5 µm was used and the number of co-localised spots was divided by the total spots counted in the analysed field of view for each image, and the average calculated across 21 images encompassing four experimental repeats.

To create the linescans for both STED and confocal data, Imaris was used to pick spots at the intended beginning and end of the line of interest. The intensity profile of this line was exported and data from at least 30 structures were used to calculate the averaged linescan via 2 degree local polynomial regression, performed using the loess function from the stats package in R. For TFG and HaloTag-Sec23A, the line originated at the highest intensity center of the TFG structure and passed through the entirety of the neighbouring HaloTag-Sec23A structure. For HaloTag-Sec23A and Sec16A, the individual linescans were generated by measuring the fluorescence originating adjacent to the Sec16A puncta and passing through the entirety of both the Sec16A and neighbouring Sec23A puncta. The intensity values were normalised for display purposes.

### EM

#### Grid preparation

We used Quantifoil R 2/2, Mo 200 mesh grids with Holey Carbon Films (Quantifoil Micro Tools GmbH) and UltrAuFoil R 2/2, Au 200 mesh grids with Holey Gold Supports (Quantifoil Micro Tools GmbH). Grids were secured in a 6-well plate using the Linkam cartridge and cartridge holder system, all of which were plasma cleaned using the Tergeo-EM (PIE scientific) and irradiated with UV light for 1 hour at room temperature to sterilize.

Grids, cartridge and cartridge holder were incubated with media for 30 mins before plating Halo-Sec23 RPE-1 cells at a density of 2.5 x 10^5^ cells/mL. The cartridge system was fully submerged with media. Cells were cultured on grids for ∼ 24 hours (37°C with 5% CO2). Cell confluency and distribution was visually inspected using a phase contrast optical microscope before staining with 1 μL/mL Oregon Green-HaloTag Ligand (Promega) following manufacturer instructions. Cells were then moved to a 37°C incubator. Each cartridge, containing a maximum of 3 grids, was moved into an individual cartridge holder and tissue culture dish, allowing batches of 3 grids to be processed for plunge freezing, minimizing the time each grid spends outside the incubator.

Grids were manually back blotted to remove media. 4 μL of 0.5 mg/mL Dynabeads MyOne Carboxylic Acid (ThermoFisher Scientific) resuspended and diluted in 10% glycerol, were added to the cell coated grid face and incubated, horizontally for ∼ 30 s at room temperature before transfer to the LeicaGP (Leica) chamber (37°C and 70% humidity). Grids were then back-blotted for 8 s before plunge freezing in liquid ethane and stored in liquid nitrogen.

Cryo-grids were clipped into autogrids (Thermo Fisher Scientific) and screened for ice thickness and cell distribution on a 200kV Glacios or Talos electron microscope (Thermo Fisher Scientific) prior to milling.

#### Cryo-FIB Lamella preparation

Lamellae were prepared on an Aquilos 2, a dual beam FIB/SEM equipped with an iFLM correlative system (Thermo Fisher Scientific) and long-operation liquid nitrogen dewar with 175 L capacity (SubAngstrom).

First, SEM tile-sets were collected and aligned to the previously acquired TEM atlases in MAPS v3.30 (Thermo Fisher Scientific). Potential areas for milling were identified based on suitable ice-thickness and their location within the grid square. Next, fluorescence Z-stacks (0.1 µm steps) were acquired at these sites using the integrated iFLM module - 20x objective (Zeiss Epiplan-Apochromat, NA 0.7, Piezo-driven), filter cube (Semrock LED-DA/FI/TR/Cy5-B-000, Quadband), camera (Basler ace 2, 2A4504-5gmPRO; Sony IMX541 CMOS sensor) and LED source (CoolLED, 365 nm/450 nm/550 nm/635 nm). The reflection, 470 nm and 565 nm channels were acquired at 0.5%, 10% and 10% laser power and 1.5 ms, 200 ms and 200 ms exposure time respectively.

The fluorescence and reflection Z-stacks were imported into MAPS and aligned to the SEM tile-set using three reference points. Clusters of fluorescent signal were preferentially chosen as sites of interest to increase the likelihood of milling through COPII-rich regions in the cell. Precise targeting of the site of interest was further aided by selection of a corresponding fluorescent Dynabead fiducial found in the same channel.

Post fluorescence imaging, grids were sputter coated with inorganic platinum (Platinum Black) for 15 seconds at 30 mA, then coated with organometallic platinum (Trimethyl(methylcyclopentadienyl)platinum(IV)) using the gas injection system (GIS) for 45 seconds, followed by another sputter coat for 15 seconds at 30 mA to further enhance sample conductivity.

Lamella positions, eucentric height and milling angles were automatically calculated in AutoTEM 2.4 (Thermo Fisher). Final lamella placement was guided by the site of interest/fiducial pair. The length and position of the lamella was optimised for each cell. Lamellae were produced using a Gallium beam, operating at 30 kV, in a stepwise manner starting at a beam current of 1.0 nA for rough milling and decreasing to 0.5 nA and 0.3 nA for medium and fine milling. Thinning was then performed in two stages with a beam current of first 50 pA, and then 30 pA. Target milling angles ranged from 9° to 15° and the target lamella thickness was set between 150 and 250 nm. Following thinning, 0.2 µm step Z stacks were acquired as described above, adjusting the 470 nm exposure time as required to visualise on-lamella signal and determine the targeting success of the milling (Extended Data Fig. 1d, right panel). Prior to unloading a second sputter coat at 7 mA for 15 seconds was applied to reduce beam-induced specimen movement of the lamella in the TEM. Grids were unloaded and then immediately loaded into a Titan Krios G2 TEM microscope (Thermofisher) to limit lamella breakage and reduce ice contamination during storage.

#### Cryo-electron tomography data collection

Montages were collected from intact lamella (3 nm pixel size with 100 um defocus) in a Krios G2 TEM running Tomography v5.19. The montages were imported into MAPS software (Thermo Fisher Scientific) where they were manually aligned with fluorescence Z-stacks.

Lower-magnification tilt series (Supplementary Table 1) with a high defocus, restricted tilt range and lower dose were collected from areas on the lamella that retained the green fluorescent signal (Supplementary Table 1). On the fly processing allowed 3D reconstructions to be quickly generated. Despite accumulating less than 10 electrons/Å^2^, the resulting tomograms were of suitable clarity and contrast to identify various intracellular structures, including membrane buds and vesicles (Fig. 1a,c). Informed by this low-magnification 3D data, we selected potential regions of interest for high-magnification data collection at 2.24 Å/pixel (Supplementary Table 1). These were chosen based on the presence of vesicular or tubular membranes in the vicinity of the endoplasmic reticulum, which was easily identified by its decoration with ribosomes (Extended Data Fig. 2c,g and Fig. 1b,d).

### Tomogram Processing

#### Tilt series alignment and reconstruction

The low magnification, low dose, restricted tilt-series were reconstructed using IMOD’s motion correction (alignframes)^65^ and AreTomo v.1.3.4^66^ for alignment and reconstruction, allowing quick access to reconstructed tomograms for visual inspection with minimal manual input. The resulting 138 tomograms were binned to a final pixel size of 14.6 Å and were inspected using IMOD. 94 tomograms were identified as containing regions indicative of ERES and were used to direct high magnification tilt series acquisition.

High-magnification tomograms were initially reconstructed as described above and IMOD was used to inspect and identify tomograms containing regions of interest. 63 tilt series containing regions of interest were reprocessed following the WarpTools pipeline, including motion correction, CTF and defocus handedness estimation (https://warpem.github.io/warp/). Tilt series were aligned using the WarpTools etomo patch tracking wrapper^65^. Bad tilts were manually removed, and poorly aligned tilt series were manually improved with etomo. CTF-corrected tomograms were reconstructed at a pixel size of 10 Å and used to pick ribosomes for further improvement of alignments with Warp/M^40^.

PyTOM_TM^67^ was used to identify ribosomes in reconstructed tomograms, using a human ribosome map (EMD-36178) as a template (Extended Data Fig. 3). Per-tomogram thresholds to select the highest cross-correlation peaks were automatically determined. Most peaks clearly overlapped with ribosomes (Extended Data Fig. 3). False positives were easily identified as these tended to cluster on dark contamination features, and were manually cleaned using the ChimeraX plugin ArtiaX^68^, obtaining a total of 16543 ribosomes. Picked ribosomes were exported at a pixel size of 4 Å, averaged and low pass filtered to 150 Å to create a starting reference, which was used to refine the ribosome dataset using relion 3D Refine^69^, achieving a resolution of ∼20 Å. To test for reference bias and confirm tomogram handedness, we repeated the relion refinement step using a mirrored copy of the starting reference. This yielded a ribosome map which had reverted to the correct hand and was in a rotated orientation compared to the template used for picking (Extended Data Fig. 3). Reassured that the ribosome structure we obtained was real, we further refined it using M, obtaining a map at ∼15 Å resolution after 5 iterations (Supplementary Table 2). The resulting alignments were used for a final round of tomogram reconstruction, yielding optimised CTF-corrected tomograms suitable for downstream analysis.

#### Tomogram analysis

Tomograms were denoised using the WarpTools denoiser prior to segmentation with membrain-seg v2^70^. Deepdewedge^71^ was also used as an alternative denoiser for tomogram visualisation and figure/movie creation. Membranes were manually segmented using Segger^72^ and individual coated membranes were initially labelled as COPI or COPII based on their visual appearance (Fig. 2).

Segmented membranes from all coated vesicles (COPI and COPII together) were expanded by 15 pixels and used as masks for template matching against non-denoised, CTF corrected tomograms using PyTOM_TM. We used EMD-19417 as template to search for COPII, and EMD-3968 to search for COPI, using particle diameter of 240 pixels, and phase flip. All peaks were extracted and manually thresholded based on their cross-correlation value using ArtiaX. Template matching confirmed our manual assignments, with the COPII template giving the highest cross-correlation peaks predominantly around membranes which were originally labelled as COPII coated, and vice versa.

Cytoskeleton was segmented in some tomograms using the Amira software (Thermo Fisher Scientific). Intermediate filaments were segmented using the AI-aided segmentation, while microtubules were segmented manually.

#### COP particle picking and alignment

Coordinates oversampling segmented membranes from the 10 Å/pixel reconstruction were defined using in-house scripts described in Pyle et al^73^. Points were separated by 50 Å and yaw and pitch angles were assigned based on particles having a normal orientation to the membrane. In plane-rotation was randomised. Local curvature was calculated based on the angle between each point and its neighbours, and points with local curvature defined by diameters smaller than 30 Å were removed. Particles from either COPII or COPI membranes were extracted in 64 voxel boxes and averaged using dynamo^74^. Particles were aligned for 5 iterations against the starting average using dynamo, allowing for no angle refinements, and shifts of 2x2x6 pixels in x, y and z direction respectively, and applying a low pass filter of 12 pixels. A shell-shaped mask generously encompassing the membrane and the coat layers was used.

#### Analysis of Ribosome exclusion zones

We used ArtiaX boundary model on ribosome coordinates with an alpha value of 0.7 to delineate a tight boundary (Extended Data Fig. 4b) defining a ribosome ‘inclusion zone’. We then created another boundary model using alpha = 1, which loosely encompasses all ribosomes (Extended Data Fig. 4c). We calculated masks from both boundaries and subtracted the volume of the first from the volume of the second to generate the volume of the ribosome exclusion zone. This allowed us to obtain an approximate value for the volume of the exclusion zone (overestimated due to the indents around the surface in the 0.7 boundary mask, and underestimated due to the fact that the exclusion zone might be thicker than the lamella). This method was used for a subset of tomograms (n = 20).

Ellipsoids encompassing ERES were fitted around points oversampling all coated membranes in each tomogram, and ribosomes were counted inside and outside exclusion zones using python scripts (https://github.com/KTDownes/Multi-scale-Molecular-Imaging-of-Human-Cells-reveals-COPI-and-COPII-Vesicles-at-ER-Exit-Sites).

#### Distance measurements

Distances were measured between clusters of COPII and COPI vesicles (Fig. 4c,d), between each COPII vesicle and its COPII or COPI neighbours (Fig. 5e-g) from coordinates of points oversampling coated membranes (https://github.com/KTDownes/Multi-scale-Molecular-Imaging-of-Human-Cells-reveals-COPI-and-COPII-Vesicles-at-ER-Exit-Sites).

#### Morphology analysis

Segmented coated membranes were manually classified as vesicles or buds based on their corresponding tomogram density. To measure circularity (Extended Data Fig. 7b), we calculated axial 2d projections of segmented vesicle membranes, and fitted an ellipse (https://github.com/KTDownes/Multi-scale-Molecular-Imaging-of-Human-Cells-reveals-COPI-and-COPII-Vesicles-at-ER-Exit-Sites). The morphology of these structures was analysed using scripts adapted from^43^ (https://github.com/KTDownes/Multi-scale-Molecular-Imaging-of-Human-Cells-reveals-COPI-and-COPII-Vesicles-at-ER-Exit-Sites). These scripts approximated the segmented membrane structure to a fitted sphere and ellipsoid. The best fitted ellipsoid and sphere for each structure was manually inspected using pyvista^75^ to determine which fit best described the structure. The volume and of the best fitted shape was plotted using seaborn^76^ and matplotlib^77^.

#### Statistics and reproducibility

Tomography workflow, from lamella preparation to tomogram reconstruction was independently repeated three times on different batches of cells, and data were pooled for analysis.

For all confocal imaging in Fig. 4a, 4e and 6a, 2 experimental repeats were conducted for the specific combination of antibodies shown.

For the experiments in Fig. 4b and 4d, there were 3 experimental repeats used for the Sec23-Sec16 distance data, and 2 experimental repeats for the Sec23-COPI distance data.

For all STED imaging, one experimental repeat was performed.

No statistical method was used to predetermine sample size. The experiments were not randomized. The Investigators were not blinded to allocation during experiments and outcome assessment. No Data points were excluded.

Statistical analyses were performed using the stats module of SciPy^78^. The difference between both volumes and sphericities were performed using the Mann-Whitney U test and the median values were quoted as the distributions were non normal. The differences between the distributions of these groups were performed using the Kolmogorov-Smirnov test.

The two-proportion Z-test was performed to assess the statistical significance of the number of filaments in ERES versus non-ERES tomograms.

## Data Availability

Raw data for all ERES tomograms are have been deposited in the EMPIAR database (EMPIAR 13270). The two representative tomograms shown in Fig. 3 have been deposited in the EMDB database (EMD-56606 and EMD-56951). The ribosome subtomogram averaging map has been deposited in the EMDB database (EMD-56963).

Source data have been provided in Source Data. All other data supporting the findings of this study are available from the corresponding author on reasonable request.

## Code availability

All code used in this paper can be found on Github (https://github.com/KTDownes/Multi-scale-Molecular-Imaging-of-Human-Cells-reveals-COPI-and-COPII-Vesicles-at-ER-Exit-Sites)

---

**Extended Data Fig. 1.**
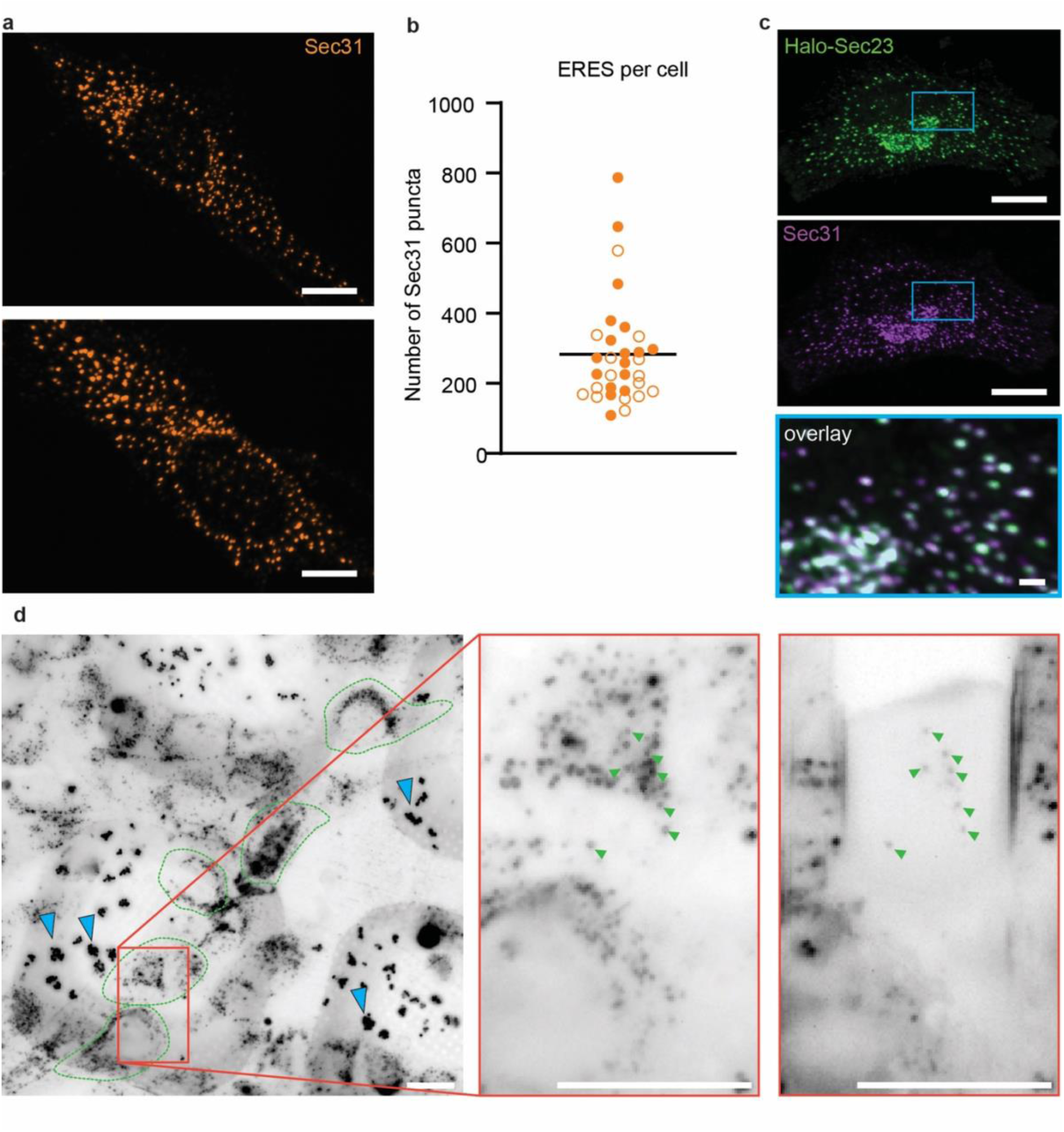
HaloTag-Sec23A is a suitable candidate for investigating ERES structure. a. Wild type RPE1 cells immunostained against Sec31A. b. Scatter plot of number of Sec31A positive structures in wild type RPE1 cells, colour coded by experimental repeat, n = 32 cells. c. HaloTag-Sec23A-RPE1 cells dyed with JFX650-HaloTag ligand (green) and co-stained using an antibody directed against Sec31A (purple). Inset highlights tight colocalisation of HaloTag-Sec23A fusion protein and endogenous Sec31A. d. Example cryo-fluorescence wide field image of HaloTag-Sec23A-RPE1 cells grown on grids stained with Oregon Green HaloTag ligand before and after milling. The punctate staining pattern throughout the cytoplasm, concentrated at the perinuclear region is compatible with Sec23 distribution, but is also contributed by autofluorescent puncta. The red box in the left panel indicates the digitally magnified region in the other panels which show cryo-fluorescence before (middle panel) and after (right panel) milling. Four cells are indicated by green dotted outlines. Blue arrow heads indicate clusters of fluorescent fiducials. Green arrows indicate fluorescent puncta visible both before and after milling. Scale bars: 10um, inset in panel c: 1um. Source numerical data are available in source data

**Extended Data Fig. 2.**
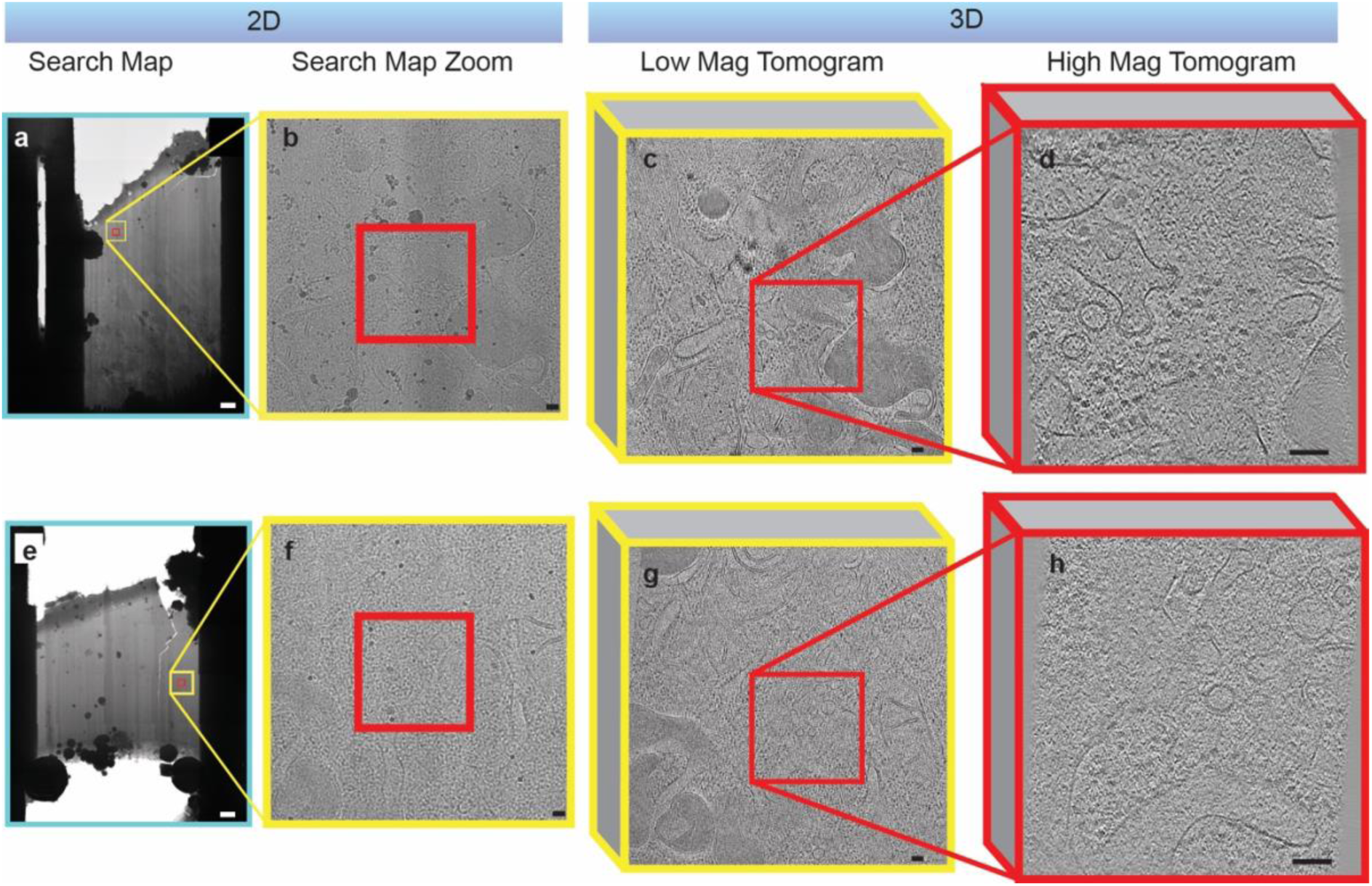
Workflow for targeting ERES on FIB/SEM lamellae. a. Lamella viewed in the TEM at low magnification (the search map). b. A digitally zoomed version of B, showing the region encompassed by the yellow box in a. The presence of ERES or more generally of membrane vesicles and tubules is not obvious. c. Slice through the xy plane of a reconstructed cryo-tomogram acquired at low magnification around the region outlined by the yellow box. d. Slice through the xy plane of a reconstructed cryo-tomogram acquired at high magnification around the region outlined by the red box. e-h. Another example as in a-e. Scale bars a,e: 1 um, b-d,f-h: 100 nm

**Extended Data Fig. 3.**
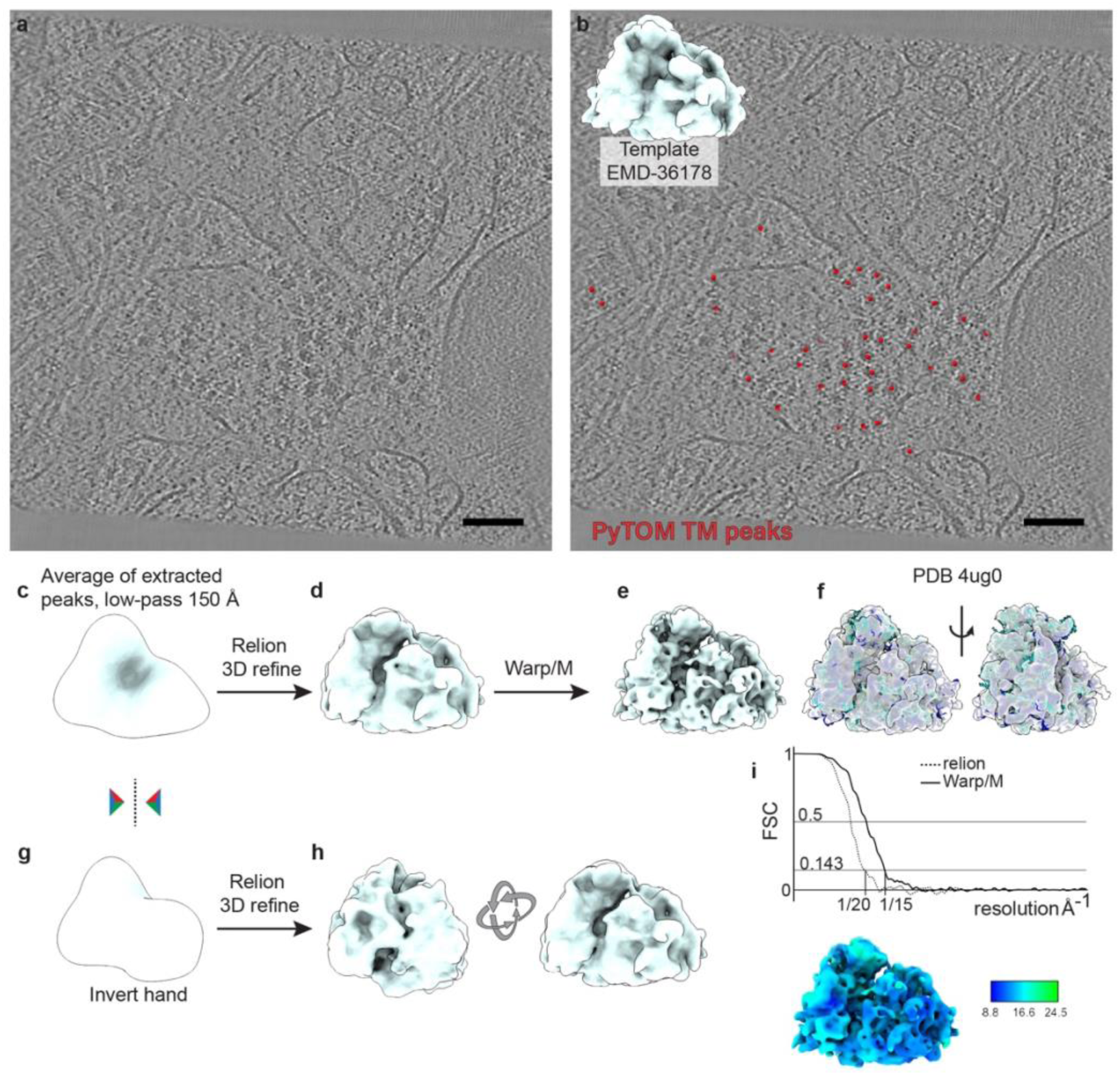
Subtomogram averaging of ribosomes. a. An xy slice through a reconstructed tomogram, where ribosome density is visible. b. As in A, with ribosome cross-correlation peaks from PyTOM_TM in red. Inset: reference used for template matching. c. Starting reference for alignments in relion, obtained by averaging PyTOM_TM peaks and low-pass filtering at 150 Å. d. The map obtained after one round of 3D refinement in relion4. e. The map obtained after 5 rounds of refinement in WARP/M. f. The map in e, with a fitted atomic model (PDB 4ug0). g. The map in c was mirrored to obtain a new staring reference for 3D refinements in relion h. The map obtained after one round of 3D refinement in relion against the mirrored starting reference, viewed in the same coordinate system (left) and rotated to match the view of the map in d. i. Top. Gold standard Fourier Shell Correlation between half maps from the 3D refinement in relion (map in panel d, dotted curve), and WARP/M (map in panel e, solid curve). Resolutions at 0.143 threshold are 20 and 15 Å respectively. Bottom. Map of ribosome coloured according to local resolution.

**Extended Data Fig. 4.**
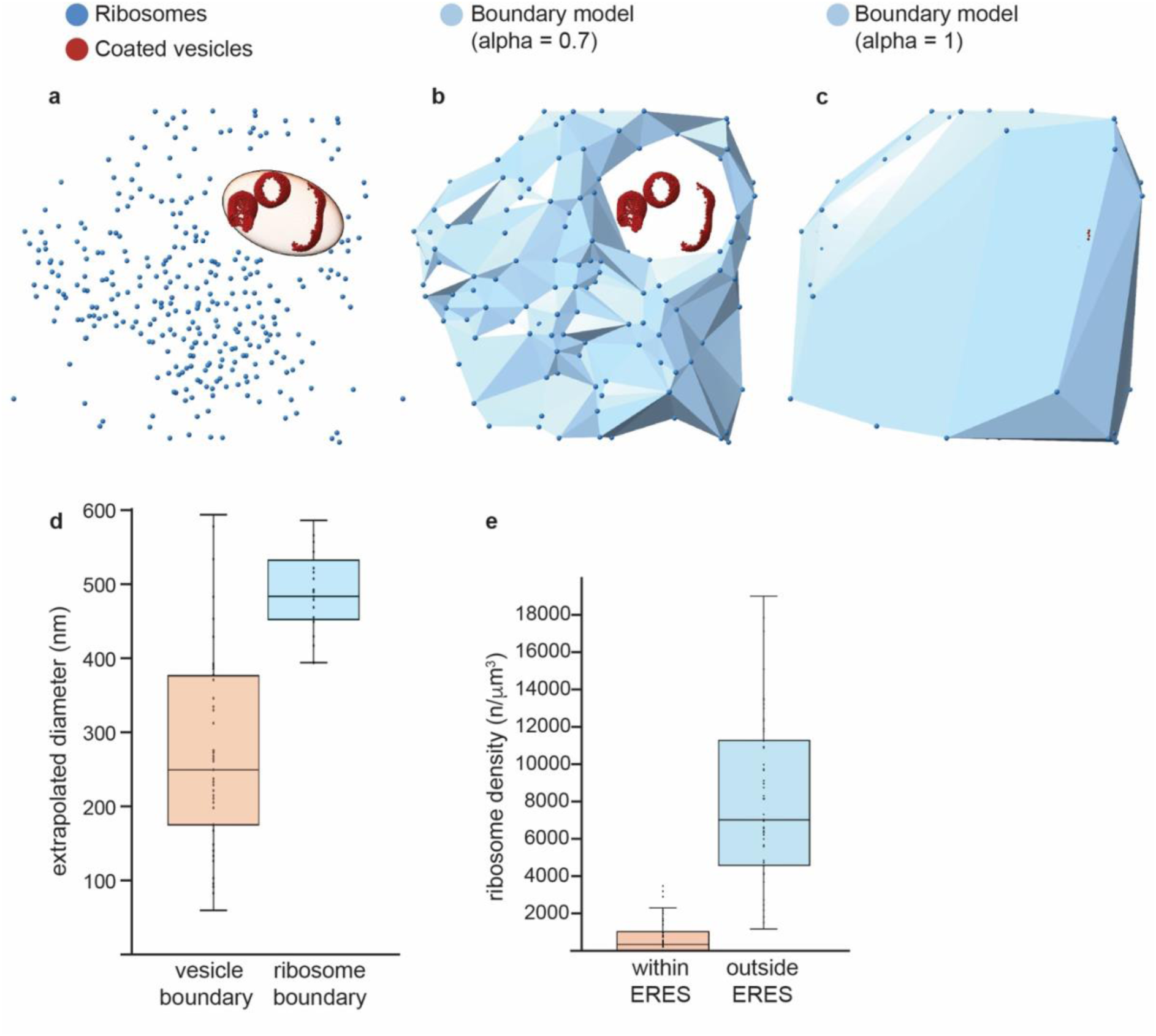
Ribosome distribution at and outside of ERES. a. Ribosome coordinates in one example tomogram as defined by template matching are shown as blue spheres. Oversamples points around coated membranes are shown in dark red. Ellipsoid fit around coated membranes to encompass ERES volume is shown in light red. b. Ribosome inclusion zone as defined in ArtiaX by setting a boundary model with alpha = 0.7 (light blue surface) c. As in b, but setting a boundary model with alpha = 1 allows to encompass the full area of the tomogram occupied by cytosol. d. Extrapolated diameters from the volumes of ERES (as from ellipsoid in a, orange), and from the volume of the ribosome exclusion zone (as from the difference between the models in c and b, light blue). e. Distribution of ribosome density inside ERES (orange) and outside the ribosome exclusion zones (light blue).

**Extended Data Fig. 5.**
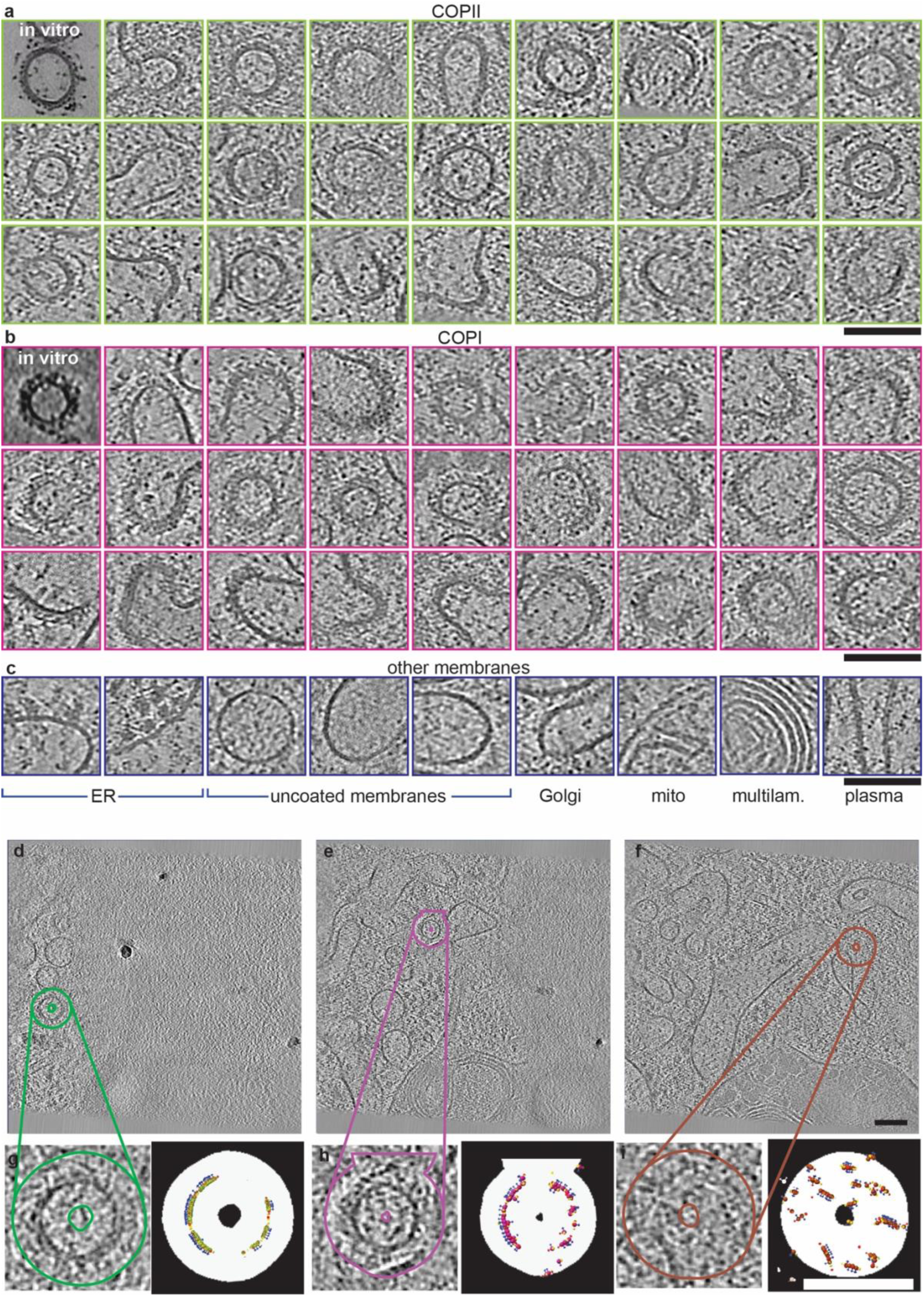
Morphology and identity of coated vesicles. a. Gallery of xy slices through COPII-coated vesicles from cryo-lamellae. Top-left panel shows an in vitro reconstituted vesicle for comparison16. b. Gallery of xy slices through COPI-coated vesicles from cryo-lamellae. Top-left panel shows an in vitro reconstituted vesicle for comparison28. c. Gallery of xy slices through other classes of membranes, as described below each panel. d. Slice through the xy plane of a tomogram overlayed with the outline of a mask used to run template matching. The mask at this plane is seen around a COPII coated vesicle. e. Slice through the xy plane of a tomogram overlayed with the outline of a mask used to run template matching. The mask at this plane is seen around a COPI coated vesicle. f. Slice through the xy plane of a tomogram overlayed with the outline of a mask used to run template matching. The mask at this plane was randomly applied as control. g. Left: a zoomed in view of a COPII vesicle from a. Right: a map of in vitro assembled COPII inner coat (EMD-19417) was used as template to search cryo-tomograms that were loosely masked around all coated membranes. Highest cross-correlation peaks are mapped onto the tomogram using ArtiaX, and a representative COPII-coated vesicles is shown. Peak positions are shown in green, and their orientations by blue, yellow and red arrows. The mask profile is in white against a black background. h. Left: a zoomed in view of a COPII vesicle from b. Right: a map of in vitro assembled COPI coat (EMD-3968) was used as template to search cryo-tomograms that were loosely masked around all coated membranes. Highest cross-correlation peaks are mapped onto the tomogram using ArtiaX, and a representative COPI-coated vesicles is shown. Peak positions are shown in purple, and their orientations by blue, yellow and red arrows. i. Left: a zoomed in view of a random region from c. Right: a map of in vitro assembled COPI coat (EMD-3968) was used as template to search cryo-tomograms that were loosely masked at random places (a mask was applied to the wrong tomogram). Highest cross-correlation peaks are mapped onto the tomogram using ArtiaX, and a representative COPI-coated vesicles is shown. Peak positions are shown in brown, and their orientations by blue, yellow and red arrows. Scale bars: 100 nm

**Extended Data Fig. 6.**
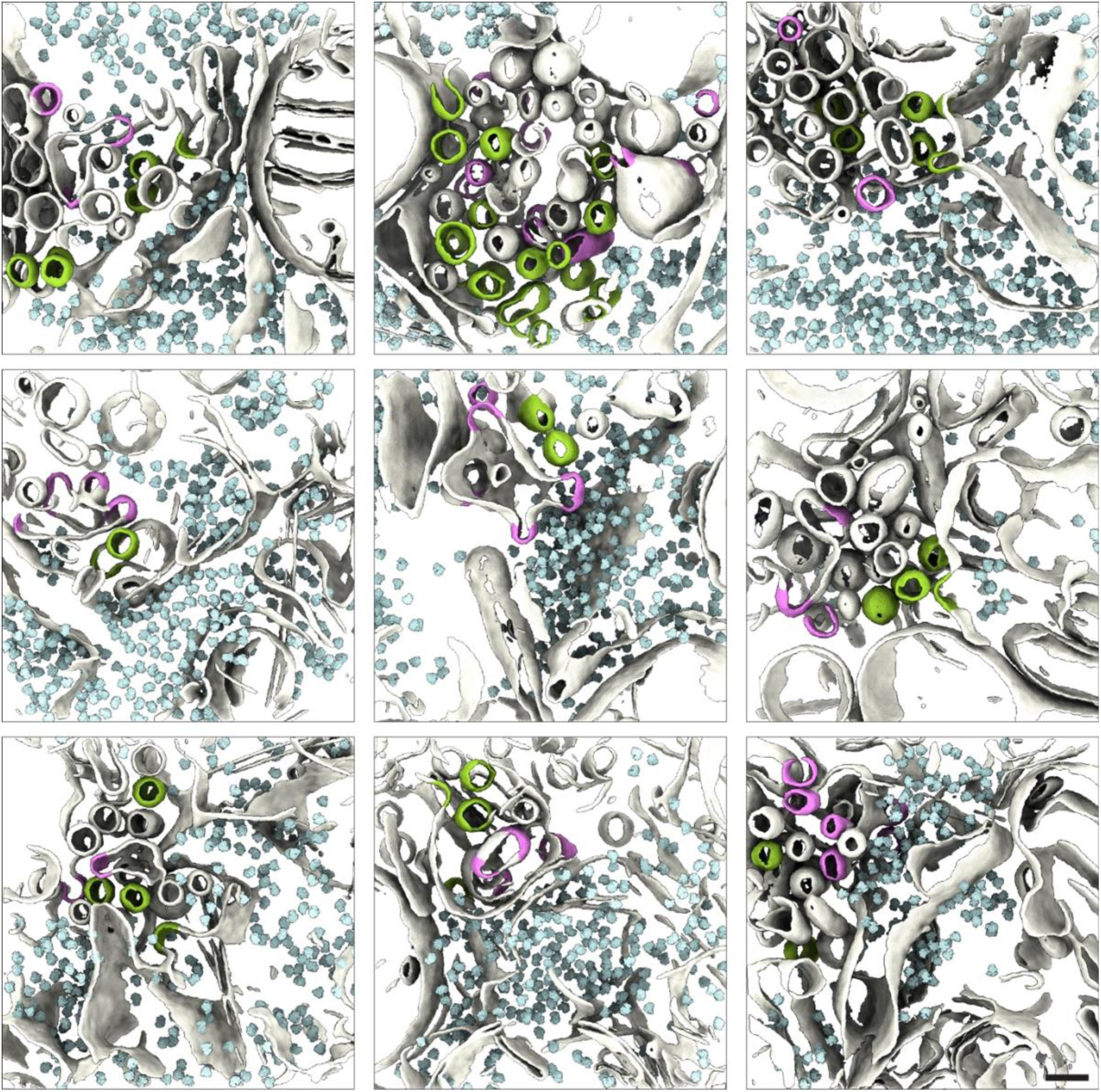
Molecular organisation of ERES. A gallery of 3D rendered segmented tomograms of ERES, where membranes are white, COPII-coated membranes are green, COPI-coated membranes are purple, and ribosomes are pale cyan. Scale bar 100 nm.

**Extended Data Fig. 7.**
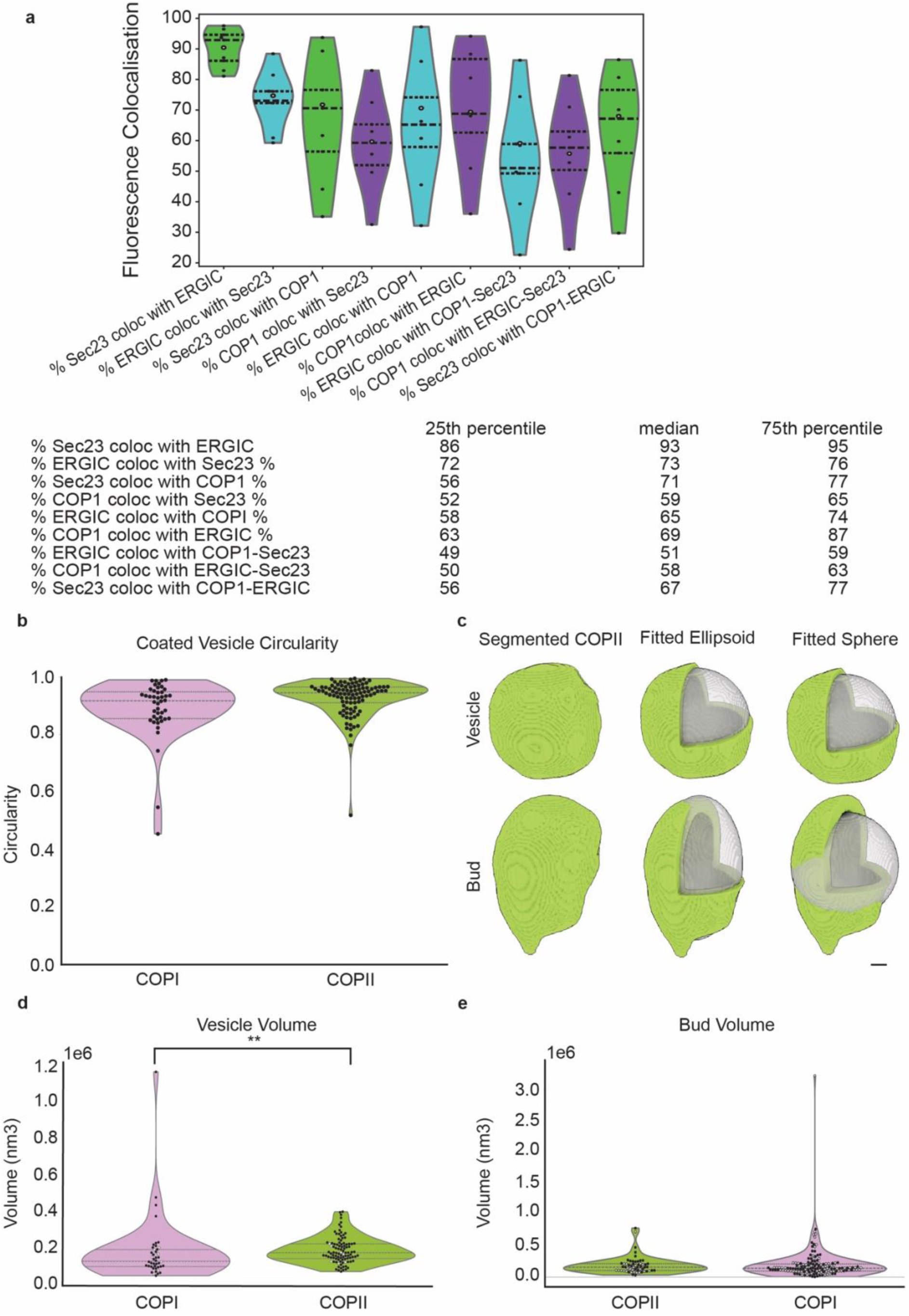
Localisation and Morphometric analysis of coated vesicles and buds. a. Quantitation of the colocalisation between fluorescent markers against COPII (Halo-Sec23A), ERGIC (ERGIC-53 antibody) and COPI (𝛽-COP antibody). Green indicates colocalisation as a percentage of COPII structures, cyan indicates colocalisation as a percentage of ERGIC structures and purple indicates colocalisation as a percentage of COPI structures. Each black point represents the percentage colocalisation reported per image, where each image contains 1-2 cells. The white points represent the percentage colocalisation reported across all images n = 9. The dashed lines represent the median and interquartile range. b. Violin plot of the circularity of COPI and COPII coated vesicles measured by fitting ellipses to axial projection of segmented vesicles. Grey dashed and dotted lines represent the median and interquartile range respectively. COPI median and IQR = 0.92, 0.86-0.95. COPII median and IQR = 0.95, 0.91-0.97. c. Examples of fitted spheres and ellipsoids for a segmented COPII vesicle (top), and bud (bottom). On the right panels, a slice from the segmented density has been cut out for clarity. Scale bar 10 nm d. Violin plot of the estimated volume of COPI and COPII coated vesicles. White and black circles indicate vesicles best fitted by a sphere and ellipsoid respectively. Grey dashed and dotted lines represent the median and interquartile range respectively. COPI median and IQR = 134,153 nm3, 105,256 – 198,030 nm3. COPII median and IQR = 180,760 nm3, 148,776 – 228,868 nm3. P=0.0014. e. Violin plot of the estimated volume of COPI and COPII coated buds. White and black circles indicate buds best fitted by a sphere and ellipsoid respectively. Grey dashed and dotted lines represent the median and interquartile range respectively. COPII median and IQR = 158,886 nm3, 106,711 – 214,387 nm3. COPI median and IQR = 141,587 nm3, 98,046 – 210,180 nm3. P = n.s. Source numerical data are available in source data

**Extended Data Fig. 8.**
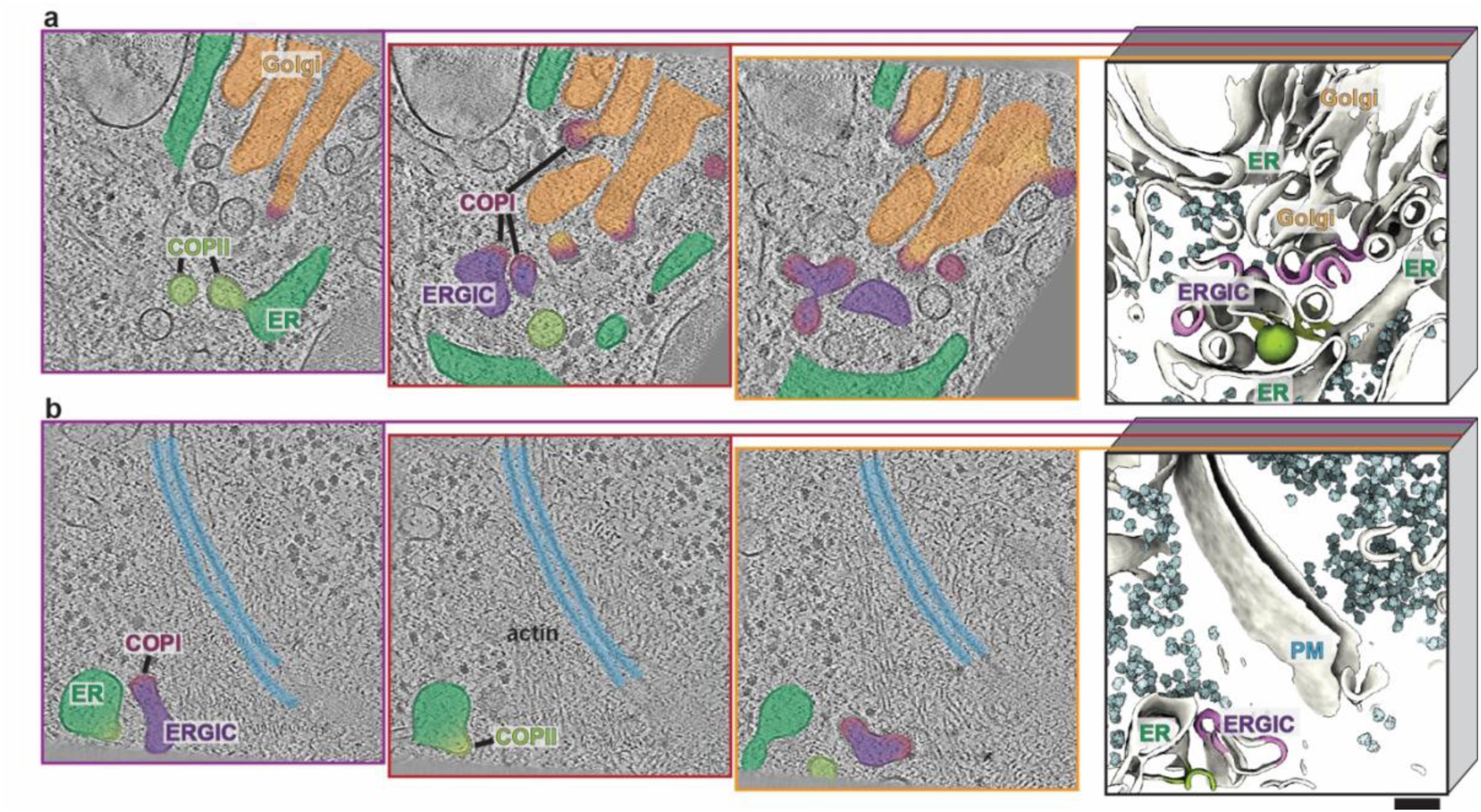
Perinuclear and peripheral ERES. a. Left: three xy slices through different z planes of a reconstructed high-magnification tomogram (3D-rendered as in Figure 3 on the right). Features have been coloured for clarity: with Golgi in orange, COPI buds and vesicles in light purple, ERGIC in dark purple, ER in dark green and COPII vesicles in light green. b. As in a, but no Golgi membranes are visible. The plasma membrane is depicted in azure. Scale bar 100 nm Source numerical data are available in source data

